# Spatially restricted *bmp16* in an extra-embryonic syncytium promotes dorsoventral patterning

**DOI:** 10.64898/2026.10.05.756835

**Authors:** Haley Couturier, Hannah Greenfeld, Jay W. Zussman, Daniel E. Wagner

## Abstract

How nuclei within a shared cytoplasm organize gene expression programs to direct developmental signaling remains incompletely understood. Here, we combine single-nucleus RNA sequencing, spatial transcriptomic mapping, and functional perturbations to characterize the zebrafish yolk syncytial layer (YSL), an extra-embryonic syncytium that supports embryonic development and patterning. We identify extensive transcriptional covariation among YSL nuclei, revealing coordinated multigene programs organized along dorsoventral and animal-vegetal axes. Within the YSL, we identify an animal-ventral subdomain expressing several BMP ligand transcripts including *bmp16, a* member of the BMP2/4 subfamily. We found *bmp16* expression to be exclusively extra-embryonic during gastrulation, and functional perturbations reveal that *bmp16* is sufficient to ventralize embryos upon overexpression and acts synergistically with *bmp2b* to promote ventral fates. Ventral YSL expression of Bmp transcripts depends on a transcriptional program downstream of *mxtx2* and is maintained independently of BMP pathway autoregulation, while negative regulation by *hhex* restricts *bmp16* expression to the ventral YSL. Together, these findings establish the YSL as a spatially organized syncytium in which coordinated nuclear gene expression programs support localized sources of embryonic patterning signals.

## Introduction

Extra-embryonic tissues are among the first to differentiate during early development and serve several critical functions in nutrient and metabolic support, morphogenesis, and axis patterning. In mammals, the anterior visceral endoderm (AVE) restricts posteriorizing Nodal and Wnt signals to establish anterior-posterior identity in the adjacent epiblast (Thomas and Beddington 1996). In birds, the hypoblast performs an analogous anterior patterning role, and in teleosts, the yolk syncytial layer (YSL) directs dorsoventral and mesendodermal patterning of the blastoderm (Bertocchini and Stern 2002; Chen and Kimelman 2000; Fan et al. 2007). Despite morphological and developmental differences, these diverse tissues share a capacity to provide signaling cues to embryos during development. However, both the molecular determinants and mechanisms by which gene expression is organized within a syncytium remain incompletely understood.

To investigate the role and nature of extra-embryonic gene expression programs in embryonic patterning, we focused on the zebrafish YSL during gastrulation. Positioned at the interface between the yolk cell and embryonic blastomeres, the YSL performs well-established functions in lipid metabolism, transport, and coordinates large scale morphogenetic movements (e.g. epiboly) during gastrulation (Carvalho and Heisenberg 2010). Expression of apolipoproteins and secretory machinery in the YSL has been shown to mediate the transfer of yolk-derived lipids to the developing embryos prior to circulatory formation around 24 hours post fertilization (hpf) (Otis et al. 2015; Marza et al. 2005; Fraher et al. 2016; Miyares, de Rezende, and Farber 2014). This metabolic function has led to frequent comparisons between the YSL and the liver, which similarly assumes similar roles lipid processing and transport later in development.

Alongside its metabolic and morphogenetic functions, the YSL is acts as a signaling center during initial embryonic axis specification. Prior to gastrulation, dorsal determinants are transported via cytoplasmic streaming from the vegetal yolk to the YSL on the future dorsal side of the embryo (Mizuno et al. 1999). By 3 hpf, nuclear β-catenin has accumulated in the dorsal YSN transducing WNT signaling and specification of dorsal organizer genes such as *dharma* (Lu et al. 2011; Fekany et al. 1999). Injection of RNase into the yolk at 3 hpf abolishes yolk transcription and subsequently causes epiboly failure illustrating that signaling from the YSL is required for development (Chen and Kimelman 2000). At approximately the same stage, maternally supplied Nanog activates the YSL transcriptome, initiating *mxtx2* transcription and downstream YSL specification (Hong et al. 2011; C. Xu et al. 2012; Gagnon, Obbad, and Schier 2018). Nodal signals emanating from the YSN are critical for mesendodermal patterning in the overlaying blastoderm, and further studies have identified YSL-derived *bmp2b* as required for ventral fate specification downstream of the transcriptional regulators Max and Mga (Sun et al. 2014; Ober and Schulte-Merker 1999; Chen and Kimelman 2000; Feldman et al. 1998; Rodaway et al. 1999; Mizuno et al. 1999).

The zebrafish blastula is comprised of three distinct lineages including the enveloping layer (EVL), the YSL, and the deep embryonic blastomeres. The YSL forms at approximately 3 hpf when marginal blastomeres deposit their contents into the underlying yolk cell forming the YSL (Kimmel and Law 1985). These yolk syncytial nuclei (YSN) undergo three to five additional rounds of division before becoming post-mitotic and migrate to form a monolayer interface between the blastomeres and yolk cell (Kane, Warga, and Kimmel 1992; Kimmel and Law 1985). By mid-gastrulation, YSN have been described as two morphogenetic domains wherein marginal YSN (external YSN, eYSN) move vegetally or undergo convergent extension while the internal YSN (iYSN) undergo convergent extension movements (Fei et al. 2019; D’Amico and Cooper 2001; Carvalho et al. 2009). However, whether eYSN and iYSN represent transcriptionally distinct states has not been fully resolved.

Several bulk profiling efforts including the use of microarrays, bulk RNA-sequencing, and classic *in situ* hybridization experiments have identified genes with YSL-enriched expression relative to embryonic cells (Hong et al. 2010; Thisse and Thisse 2008; Gagnon et al. 2018). However, most recent scRNA-seq atlases contain few or no YSN, likely owing to its syncytial nature, technical barriers to recovery following dissociation, and small population size (Wagner et al. 2018; Kukreja et al. 2024; Farrell et al. 2018; Kamimoto et al. 2023; Spanjaard et al. 2018; Sur et al. 2023; Lange et al. 2024). As a result, despite extensive single-cell profiling of the zebrafish embryo the YSN has evaded thorough transcriptional profiling at the single-nucleus level.

In this study, we leverage single nucleus RNA-sequencing (snRNA-seq) to perform, to our knowledge, the first targeted transcriptional profiling of the zebrafish YSL during gastrulation. We establish that the YSN exhibit considerable transcriptional heterogeneity and comprise several coordinated multigene programs organized along the dorsoventral and anterior-posterior axes. We identify a ventrally enriched, YSL-specific Bmp ligand, *bmp16*, that promotes embryonic ventralization. Our findings further indicate that ventral Bmp ligand expression in the YSN is regulated by a YSL-specific gene regulatory program distinct from the well-established BMP autoregulatory mechanism observed in embryonic tissues.

## Results

### Single-nucleus transcriptional profiling of yolk-associated tissues reveals yolk syncytial layer specific features

To comprehensively investigate transcriptional features and heterogeneity of the zebrafish yolk syncytial layer (YSL), we developed a strategy to enrich yolk-associated nuclei from gastrula-stage embryos for single-nucleus RNA-seq (snRNA-seq). We found that a combination of surgical enrichment, mild fixation, and single-nucleus isolation preserved yolk syncytial nuclei (YSN) for transcriptomic profiling (see Methods). We conducted this workflow on over 2,000 micro-dissected embryos at late shield stage (∼6-7 hours post fertilization (hpf); **Fig. 1A**). This developmental window shortly follows the establishment of the primary embryonic axes and encompasses a period during which the YSL acquires a domed morphology with distinguishable internal (iYSL) and external (eYSL) spatial compartments.

**Fig. 1.**
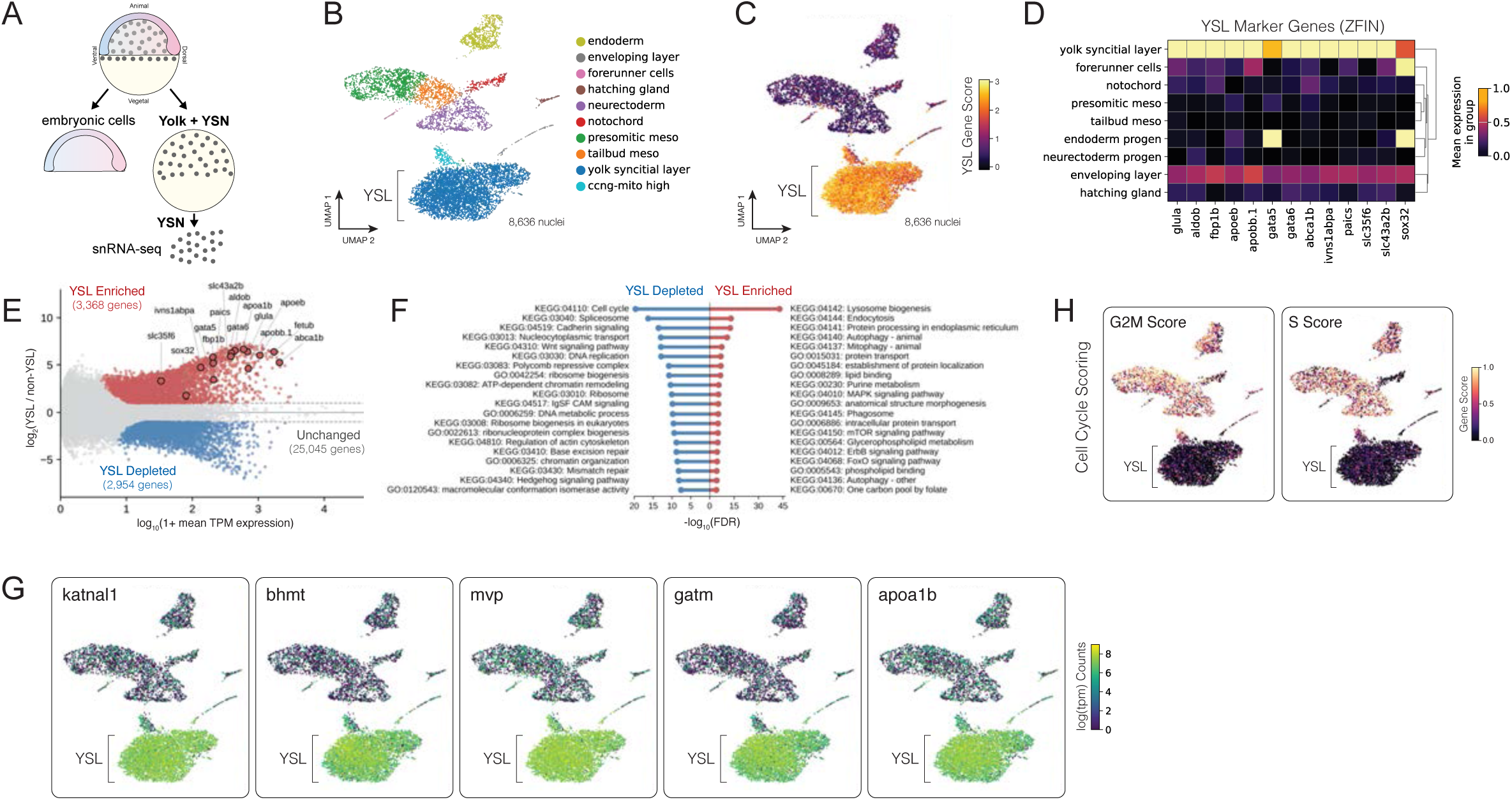
snRNA-seq transcriptomic profiling of the zebrafish yolk syncytial layer. **(A)** Experimental schematic of embryonic dissection strategy to isolate YSN for snRNA- seq. **(B)** Uniform Manifold Approximation and Projection (UMAP) embedding for 8,636 snRNA-seq profiles with cell type/tissue annotations indicated. **(C)** UMAP embedding of all nuclei colored by aggregate expression score calculated from a YSL enriched 271-marker gene set (Xu et al 2012). **(D)** Normalized expression heatmap of top YSL marker genes defined by ZFIN vs annotated groups. **(E)** Scatterplot of differential gene expression statistics for all genes: log2 fold change enrichment (YSL vs non-YSL clusters) and log10 tpm mean expression. Genes with statistically significant changes (>1 log2 fold change; FDR-adjusted p-value <0.05) are highlighted for both enriched and depleted transcripts. tpm, transcripts per million. See also Table S1. **(F)** Top 10 enriched GO/KEGG terms associated with YSL-enriched and YSL-depleted transcripts, respectively, sorted by FDR-adjusted p-value (log10). See also Table S2. **(G)** UMAP embedding of all nuclei colored by log(tpm) transcript counts for top YSL- enriched transcripts. tpm, transcripts-per-million **(H)** UMAP embedding of all nuclei colored by cell cycle scores derived from G2M-phase and S-phase associated transcripts.

Following enrichment, we performed split-pool barcoding and sequencing to generate a snRNA-seq dataset which, after quality control and preprocessing, comprised 8,636 nuclei. We next performed standard normalization, dimensionality reduction, and graph-based clustering to identify major transcriptional populations. Clusters were annotated using a combination of reference mapping to the Zebrafish Meta Atlas Project (ZMAP) (Aponte-Santiago, Su, and Wagner 2026) (**Fig. 1B, S1A-C**), expression-based scoring using a public YSL reference gene set (**Fig. 1C)** (C. Xu et al. 2012) and manual inspection of differentially expressed genes (**Fig. S1D, Table S1**). These analyses identified a prominent YSL cluster, alongside cell type clusters derived from margin- or yolk-associated embryonic tissues including mesoderm, tailbud, posterior ectoderm, endoderm, forerunner cells, and enveloping layer (**Fig. 1B, S1B-C**). The identity of this YSL cluster was further confirmed using a panel of YSL marker genes annotated by the Zebrafish Information Network (ZFIN) (**Fig. 1D**; (Bradford et al. 2022)). We estimate that our YSL cluster (n=4823 YSN) represents an approximate 18X single-nucleus sampling depth of the endogenous syncytium based on empirical YSN per embryo counts from yolk injected SYTOX labeling of YSN (**Fig. S1E-F**; ∼266 YSN per embryo).

Initial ZMAP-based automated annotations (**Fig. S1A-C**) indicated that YSN were strongly associated with liver/hepatocyte-like transcriptional signatures, consistent with known gene expression similarities between these tissues (Miyares, de Rezende, and Farber 2014). To generate a comprehensive transcriptomic profile of the YSL at 7 hpf, we performed differential expression analysis and identified transcripts that were either enriched (n=3,368) or depleted (n=2,954) relative to all other cell type clusters in our dataset (**Fig. 1E**). These YSL-enriched transcripts included all members of the ZFIN-defined marker panel (**Fig. 1D**) and were enriched for GO/KEGG terms associated with liver-like functions, e.g. vesicle trafficking, metabolism, and protein / lipid processing (**Fig. 1F, Table S2**). Top YSL-enriched genes included factors associated with methionine metabolism (*bhmt*), creatine biosynthesis (*gatm*), and lipid transport (*apoa1b*), as well as the microtubule regulator *katnal1* and major vault protein *mvp* (**Fig. 1G**).

The YSL completes its nuclear divisions by the onset of epiboly (4-5 hpf) (Veil et al. 2018; Kimmel and Law 1985; D’Amico and Cooper 2001; Kane, Warga, and Kimmel 1992), so we thus examined transcriptional patterns associated with cell cycle progression. Of note, GO/KEGG terms associated with cell cycle, DNA replication, and chromatin organization were among the most significant for YSL-depleted transcripts relative to embryonic cell types (**Fig. 1F, Table S2**). Consistent with this result, transcriptional markers of S phase (*pola1*) and G2M phase (*top2a*) were broadly expressed across embryonic clusters, but largely absent from the YSL (**Fig. S1G**). Automated cell cycle scoring confirmed these trends (**Fig. 1H**) and predicted the largest fraction of G1/G0 states among YSN (**Fig. S1H**). These data thus confirm that at 7 hpf, YSN exhibit transcriptional features consistent with a post-mitotic state. Collectively, these analyses confirm the identity of the late shield-stage YSL and provide an enriched resource for defining the YSL transcriptional landscape.

### YSL nuclei comprise orthogonal dimensions of transcriptional heterogeneity

To investigate whether distinct subsets of nuclei exist within the YSL, we first tested for the presence of transcriptional heterogeneity. To accomplish this, we subsetted the nuclei within our identified YSL cluster and performed targeted re-embedding and subcluster analyses (**Fig. 2A**). Graph-based Leiden clustering on this subset population revealed four groups with distinct transcriptional markers (**Fig. 2B, Table S3**). To corroborate this finding, we implemented non-negative matrix factorization (cNMF) to identify modules of transcriptional covariation in a cluster-agnostic manner. Joint analysis of NMF factorization stability and Frobenius reconstruction error revealed multiple values of k associated with highly stable solutions (**Fig. S2A**). We report reconstructions for both 4- and 8-component factorizations (**Fig. 2B, S2B, Table S4**). The 4-component factorization, for which the number of coordinated gene expression programs (GEPs) matched the number of Leiden subclusters, also revealed similar cell subsets and identified many of the same transcripts (**Fig. 2B**, **Table S4**). These analyses thus demonstrated that independent methods (graph clustering and matrix factorization) converged on similar signatures of transcriptional covariation within the YSL.

**Fig. 2.**
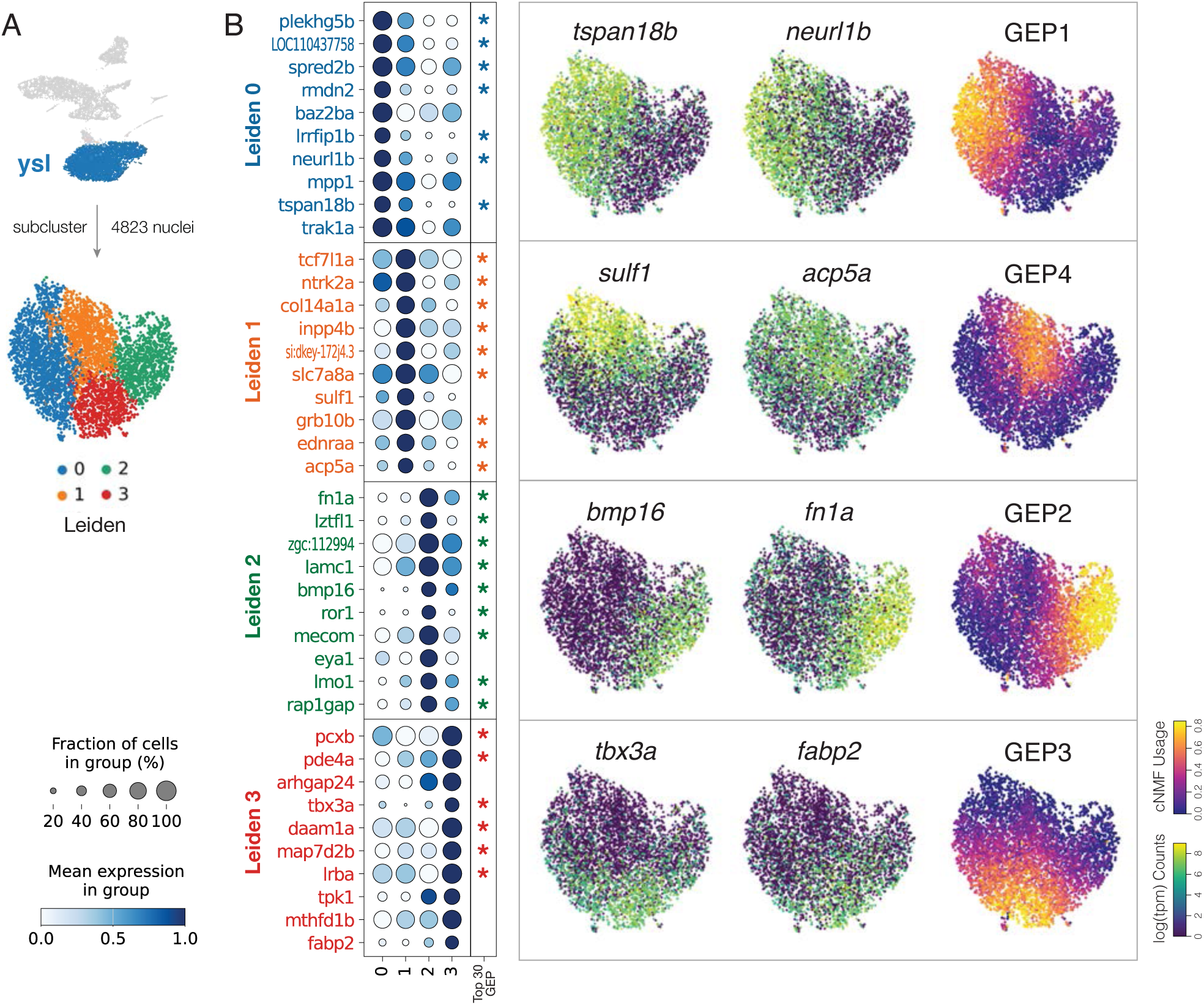
Transcriptional heterogeneity within the yolk syncytial layer. **(A)** Subset strategy, UMAP embedding, and Leiden clustering analysis of the annotated YSL population. **(B)** Left, dotplot depicting marker transcript expression vs YSL subcluster. Top 10 enriched transcripts for each subcluster are shown. Dot color and size indicate expression level and fraction of expressing nuclei, respectively. Asterisks indicate gene co-membership in the corresponding cNMF-defined components. Right, UMAP overlays of the YSL subset, colored by select marker genes and cNMF GEP scores (4- component factorization).

To investigate whether the identified GEPs correspond to spatially distinct (as opposed to intermingled or non-spatially defined) subsets of nuclei, we leveraged a public spatial transcriptomics dataset collected from zebrafish gastrula (Wan et al. 2026). This dataset included embryos collected at mid gastrulation (75% epiboly; similar to the 7 hpf collection timepoint in the present study), used probes for 495 selected developmental transcripts, and contained 892 annotated YSN. We first generated a batch-corrected Symphony-based embedding of this dataset (**Figs. S3A-B**), which recapitulated reported spatial and transcriptional covariation patterns (**Figs. S3C-D**). We next projected our snRNA-seq dataset into this Symphony reference embedding and performed kNN regression-based transfer of spatial coordinates (**Fig. 3A**, see Methods). For this analysis, we utilized phi and theta polar projection coordinates which correspond to the approximate dorsal-lateral-ventral and animal-vegetal (AV) axes of the embryo, respectively. To benchmark this approach, we first assessed spatial projections for our entire 8,626-nucleus dataset (**Fig. S3E-F)**, which recovered well-established relationships for key embryonic cell populations, e.g. dorsal anterior and dorsal posterior projections for hatching gland and notochord, respectively, and posterior/vegetal projections for tailbud/margin (**Fig. S3G-H)**. This projection strategy thus facilitates the mapping of spatial information to whole-transcriptome data without the need for imputation.

**Fig. 3.**
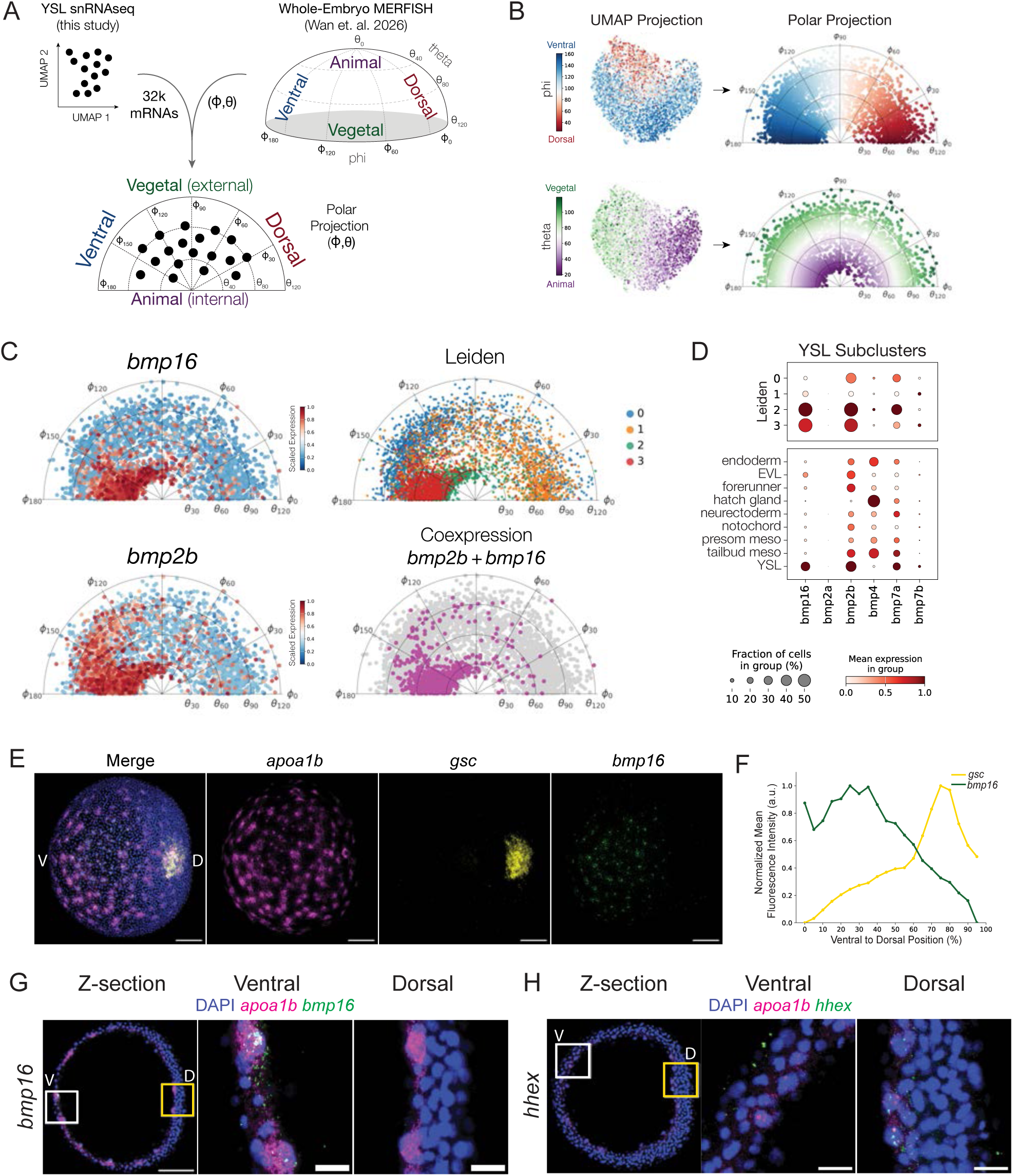
Spatial mapping of yolk syncytial layer populations. **(A)** Schematic depicting the projection of YSL nuclei into spatial polar coordinate space. **(B)** YSL polar coordinate projections for phi and theta. Paired UMAP projections for each coordinate are shown. **(C)** YSL polar coordinate projections for *bmp16, bmp2b*, Leiden subclusters, and *bmp16-bmp2b* co-expressing nuclei. **(D)** Dotplot depicting expression patterns of Bmp ligand transcripts amongst Leiden subclusters (top) and annotated tissues (bottom). Dot color and size indicate expression level and fraction of expressing nuclei, respectively. **(E)** Representative image of whole mount *bmp16* RNA FISH. YSL marker (*apoa1b*), dorsal marker (*goosecoid*; *gsc*), and nuclear marker (DAPI). Dorsal is to the right. Scale bar, 100 µm. **(F)** Quantification of YSN normalized *bmp16* and *gsc* mean fluorescence intensity across DV axis in wild-type 8 hpf embryos. (n = 21) **(G)** Representative image of 6 µm Z-stack of *bmp16* RNA FISH. Insets of ventral (white) and dorsal (yellow) regions (middle and right panels, respectively) to illustrate YSN versus embryonic cells. Scale bar, 20 µm. **(H)** Representative image of 6 µm Z-stack of *hhex* RNA FISH. Insets of ventral (white) and dorsal (yellow) regions (middle and right panels, respectively) to illustrate YSN versus embryonic cells. Scale bar, 20 µm.

We next assessed direct spatial projections between our subsetted YSN snRNA-seq profiles and the YSN component of this spatial reference. Both phi and theta polar projection coordinates mapped to our YSN snRNA-seq embedding in a non-random fashion (**Fig. 3B**). This structured mapping suggested that transcriptional subsets identified by clustering and cNMF might correspond to spatially defined regions of the YSL. Past studies have suggested a limited number of regionally expressed marker genes within the YSL, consistent with morphologically defined eYSL and iYSL regions (Hong et al. 2010). To benchmark our YSL spatial embedding, we thus examined the theta distributions of four key marker genes: *gata3* and *znf503*, reported to be expressed in eYSL and *aplnra* and *hyou1*, reported to be expressed in non-eYSL (B. Tucker et al. 2007; Hong et al. 2010; Thisse and Thisse 2008). Indeed, these four transcripts displayed predicted enrichments in vegetal/eYSL and animal/iYSL regions in our polar YSL embeddings, respectively (**Fig. S3I**). To test whether dorsoventral (phi) projections similarly recapitulate spatial relationships, we examined the expression of *hhex1*, a dorsal YSL determinant that is induced by Nieuwkoop-derived Wnt signaling downstream of *dharma* (Bischof and Driever 2004; Ho et al. 1999). Phi projections revealed moderate dorsal YSL enrichment of the *hhex1* transcript, consistent with this expectation (**Fig. S3I**). Collectively, these patterns confirm that spatial projections accurately reconstruct spatial relationships amongst transcriptionally distinct YSN.

We next examined the spatial projections of YSL subpopulations. We noted that Leiden subcluster 2 was marked by enriched expression of both *bmp16* and *bmp2b* (**Fig. 2B, Table S3**) and was predicted by spatial projections to occupy ventral YSL (**Fig. 3C, S3J**). In zebrafish embryos, Bmp ligands are expressed ventrally and promote ventrolateral fates (Mullins et al. 1996; Nguyen et al. 1998). While the expression domain of *bmp2b* has been reported to extend into ventral extra-embryonic regions, *bmp16* has been studied primarily at later developmental stages in the context of cardiac and vascular development with no prior known associations to the YSL (M. Xu et al. 2026; Feiner et al. 2009). We thus asked whether *bmp16* and *bmp2b* were regionally co-expressed in the gastrula stage YSL. This analysis confirmed that *bmp2b* was expressed broadly throughout the animal-vegetal ventral YSN, whereas *bmp16* was restricted to animal ventral regions and frequently co-expressed with *bmp2b* (**Fig. 3C**).

To illustrate the broader embryonic and extra-embryonic expression patterns of BMP family ligands, we mapped their transcriptional counts to both our ZMAP-defined cell clusters and the four YSL-defined subclusters. This analysis revealed *bmp16* expression to be uniquely YSL-specific at this stage of development, whereas both *bmp2b* and *bmp7a* expression spanned both embryonic cell types and the YSL (**Fig. 3D**). Together these data indicate that despite its syncytial nature, individual YSN display a range of structured transcriptional covariation. These identities are predicted to span at least four coordinate gene expression programs mapping onto the animal-vegetal (AV) and dorsoventral (DV) axes of the syncytium, including ventral YSL which express numerous Bmp ligands and a specific animal-ventral region that expresses a YSL-specific Bmp ligand, *bmp16*.

### *bmp16* is expressed in the ventral-anterior YSL

To validate endogenous expression patterns inferred from scRNA-seq and spatial projections, we performed whole mount RNA fluorescent *in situ* hybridization (RNA FISH) for *bmp16* together with dorsal marker, *goosecoid*, and pan-YSN marker, *apoa1b*, at 8 hpf (Otis et al. 2015). Transcripts for *bmp16* exhibited an animal-ventral distribution and were notably excluded from dorsal regions, consistent with a ventral localization predicted by our spatial transcriptomics mapping (**Fig. 3E**). To quantify this expression domain, we developed a nuclei segmentation workflow (see Methods) to distinguish embryonic nuclei from YSN and quantified RNA FISH signal intensity per nucleus. This quantification confirmed that *bmp16* expression was enriched in ventral YSN and spatially segregated from dorsal *goosecoid* expression (**Fig. 3F**). We further confirmed the ventral enrichment of *bmp16* transcripts within individual YSN at high magnification (**Figs. 3G, S3K**). In parallel, we found that expression of *hhex*, a previously described YSL dorsal determinant, was localized to dorsal YSN (**Fig. 3H, S3L**), consistent with its enriched expression in YSN subcluster 1 (**Fig. S3I**, **Table S3;** Bischof and Driever 2004).

To contextualize the expression of *bmp16* in the YSN, we asked if *bmp16* is expressed in the embryo at later developmental stages. We leveraged ZMAP to dissect the expression of *bmp16* in different cell types from 4 to 120 hpf (**Figs. S3M-N**). This analysis revealed that *bmp16* is expressed in embryonic cells starting around 14 hpf in endodermal and mesodermal tissues with highest expression levels in endothelial, heart, and intestinal cells, consistent with recent reports (Feiner et al. 2009; M. Xu et al. 2026). Expression of *bmp16* increases in these tissues around 36-48 hpf. Together, these data indicate that during early embryonic development, *bmp16* expression remains largely restricted to the extraembryonic animal-ventral YSL. This restriction is subsequently lost as *bmp16* expression expands in embryonic tissues, most prominently in endothelial and cardiac lineages.

### *bmp16* is sufficient to promote embryonic ventralization

BMP signaling is required for specification of ventral cell fates during dorsoventral patterning, and *bmp2b* and *bmp7* are required for establishing the BMP signaling gradient in zebrafish (Schmid et al. 2000; Dick et al. 2000). Global loss of function *bmp2b* and *bmp7* mutants display dorsalized phenotypes with progressive loss of the ventral tailfin, whereas overactivation of BMP signaling activity causes ventralized phenotypes displaying expanded somites and a loss of anterior neural structures (Mullins et al. 1996; Kishimoto et al. 1997; Nguyen et al. 1998).

Notably, loss of YSL *bmp2b* has been reported to result in mild dorsalizations (Sun et al. 2014). *bmp16* belongs to the BMP2/4 subfamily and has been shown to activate BMP responsive elements and canonical target genes, although less potently than *bmp2b* or *bmp4* (Marques et al. 2016; Feiner et al. 2009). Because *bmp16* is expressed in a ventral YSL domain overlapping in part with YSN *bmp2b* expression as our spatial projections suggest, we asked whether *bmp16* possesses ventralizing activity during dorsoventral patterning.

To assess BMP signaling activity, we synthesized and microinjected mRNAs for *bmp2b*, *bmp16*, and mCherry2 into one-cell stage embryos. Using previously established morphological criteria for dorsoventral patterning defects in zebrafish, embryos were phenotypically scored at 24 hpf (see Methods). As expected, *bmp2b* overexpression produced ventralized embryos ranging from mild (V1) to severe (V4) phenotypes (**Figs. 4A-B**). Strikingly, *bmp16* overexpression also induced ventralized phenotypes similar to those observed upon *bmp2b* overexpression (**Figs. 4A-B**).

**Fig. 4.**
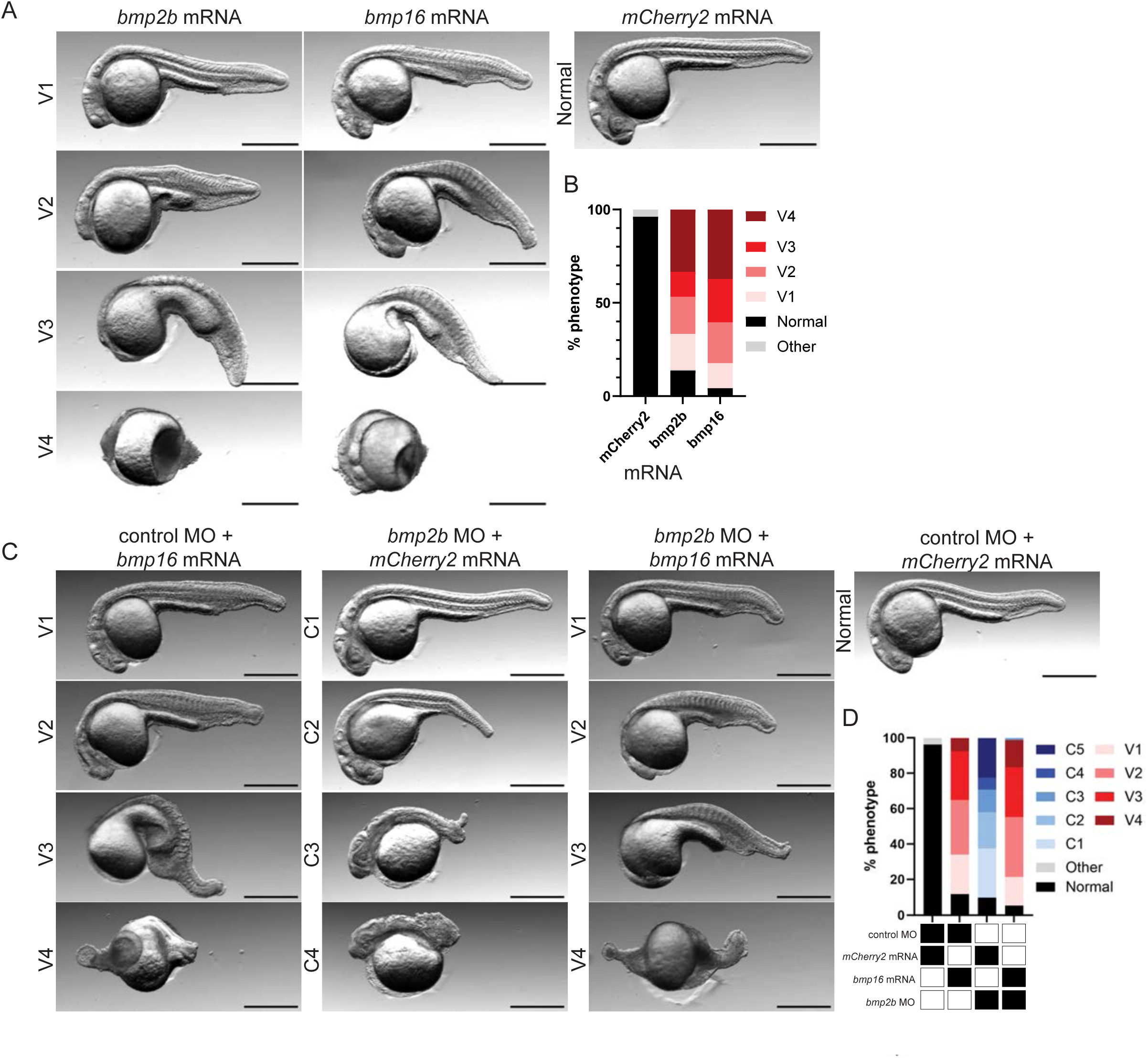
*bmp16* is sufficient to ventralize embryos. **(A)** Representative image of 5 pg/nL *bmp2b* mRNA (left panel), *bmp16* mRNA (middle panel), and mCherry2 mRNA (right panel) injected embryos at 24 hpf. Scale bar = 500 µm. (n = 246, 164, 155) **(B)** Stacked bar plot of percentage of ventralized phenotypes in mCherry2, *bmp2b*, and *bmp16* mRNA one-cell injected embryos at 24 hpf from 3 independent experiments. **(C)** Representative images of 5 pg/nL (mRNA) and 0.25 pg/nL (MO) one-cell stage injected embryos. control MO + *bmp16* mRNA (left panel), *bmp2b* MO + mCherry2 mRNA (middle-left), *bmp2b* MO + *bmp16* mRNA (middle-right) and *control* MO + mCherry2 mRNA. Scale bar, 500 µm. (n = 171, 195, 235, 197, respectively) **(D)** Stacked bar plot of dorsoventral phenotypes at 24 hpf of embryos injected with control MO + mCherry2 mRNA, control MO + *bmp16* mRNA, *bmp2b* MO + mCherry2 mRNA, or *bmp2b* MO + *bmp16* mRNA.

We next asked whether *bmp16* could functionally compensate for *bmp2b*. To determine if expression of *bmp16* could rescue the reduction of *bmp2b*, we knocked-down *bmp2b* with microinjection of a *bmp2b* morpholino oligonucleotide (MO) and co-injected *bmp16* mRNA. We microinjected embryos with combinations of MO and mRNA generating four experimental conditions: (1) control MO and mCherry2 mRNA, (2) control MO and *bmp16* mRNA, (3) *bmp2b* MO and mCherry2 mRNA, and (4) *bmp2b* MO and *bmp16* mRNA. Embryos were then scored for dorsoventral phenotypes at 24 hpf. Embryos injected with mCherry2 mRNA and control MO developed normally, while *bmp16* overexpression with control MO produced ventralized phenotypes (V1-V4) (**Figs. 4C-D**). To produce a range of dorsalizations, we titrated the *bmp2b* MO and identified a concentration where *bmp2b* knockdown result in dorsalized phenotypes ranging from mild (C1) to severe (C5). Strikingly, co-injection of *bmp16* mRNA with *bmp2b* MO suppressed dorsalization and resulted in ventralized embryos (V1-V5) (**Figs. 4C-D**). Together, these findings demonstrate that ectopic *bmp16* expression is sufficient to promote ventralization even in the context of reduced *bmp2b* activity. Based on these data, we establish that *bmp16* can function similarly to other BMP2/4 subfamily ligands during early embryonic dorsoventral patterning.

### *bmp16* acts redundantly with *bmp2b* during dorsoventral patterning

We next sought to determine the endogenous role of *bmp16* during dorsoventral patterning. We investigated the effects of *bmp16* knockdown by injecting a translation-blocking *bmp16* MO at the one-cell stage and assessed phenotypic effects at 48 hpf. *bmp16* knockdown consistently and reproducibly produced yolk-related morphological abnormalities that first appeared at the onset of yolk extension formation around 18 hpf (**Figs. S5A-D**). We quantified the yolk extension abnormalities at 48 hpf identifying a narrowed and shortened yolk extension (**Figs. S5B-C**) as well as a shortened body axis (**Figs. S5D**). Similar phenotypes were observed at 24 hpf following injection of an independent (splice blocking) *bmp16* MO, supporting the specificity of this phenotype (**Figs. 5SE-H**). However, because *bmp16* is expressed in additional cell types at later developmental stages such as cardiac and vasculature tissues, these phenotypes may reflect a combination of functions in these tissues and/or the YSL.

**Fig. 5.**
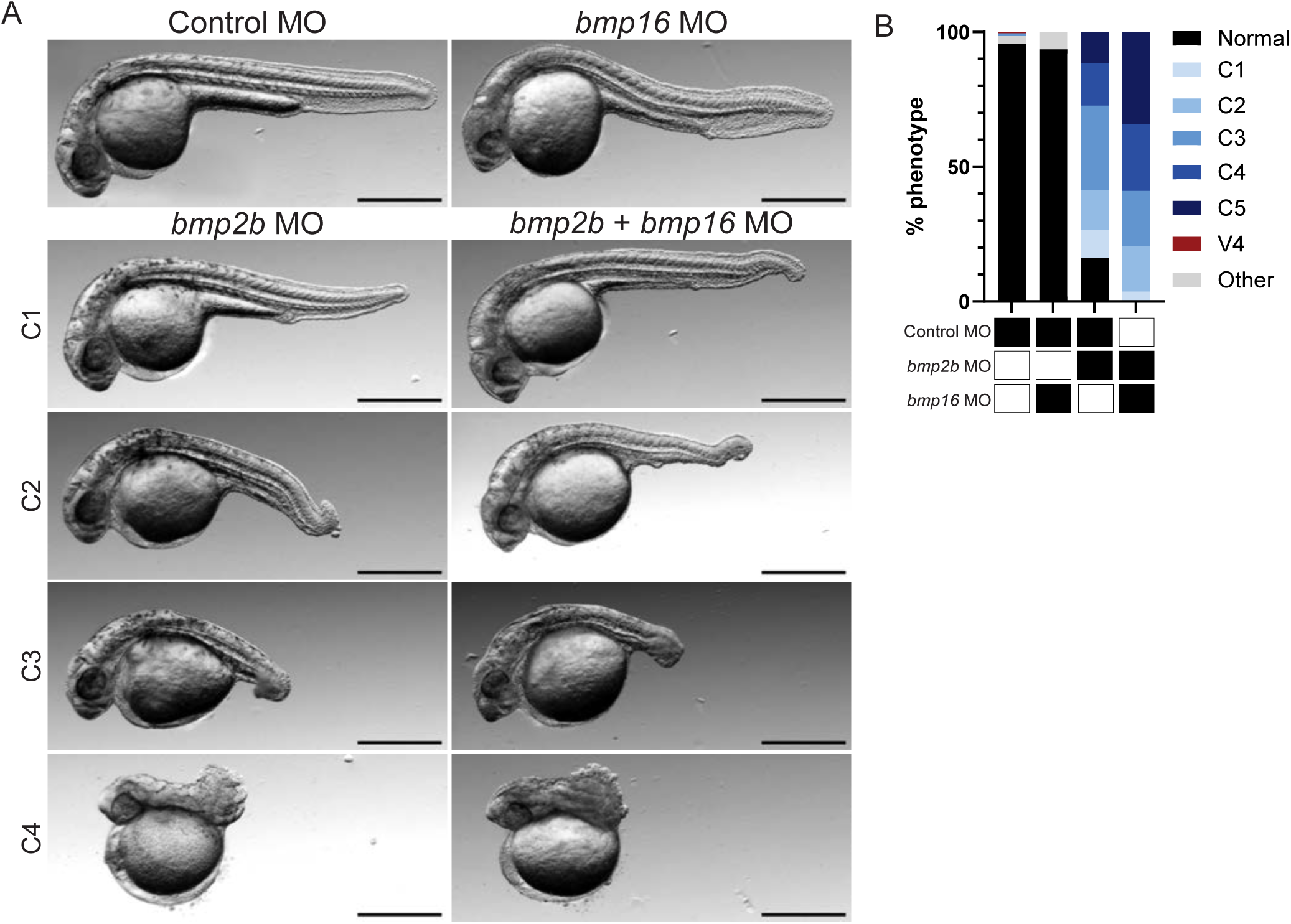
*bmp16* acts redundantly with *bmp2b* during dorsoventral patterning. **(A)** Representative images of phenotypes from control MO (top left panel) and *bmp16* MO (top right panel), *bmp2b* MO (bottom left panels), and *bmp2b* MO + *bmp16* MO (bottom right panels) one-cell injected embryos at 24 hpf. Dorsalized phenotypes are categorized from mild (C1) to severe (C4). Scale bar, 500 µm. (n = 206, 157, 227, 190) **(B)** Stacked bar plot of dorsalized phenotypes from all conditions at 24 hpf (*bmp2b* MO phenotype percentages: 16% Normal, 10% C1, 15% C2, 31% C3, 16% C4, 11% C5; *bmp2b* MO + *bmp16* MO phenotype percentages: 0% Normal, 4% C1, 17% C2, 20% C3, 25% C4, 34% C5). Fisher’s Exact Test performed between *bmp2b* MO vs *bmp16* MO/*bmp2b* MO and grouped as the following: C1 vs. Rest: p < 0.0037, **; C1+C2 vs Rest: p <0.0444, **; C1+C2+C3 vs Rest: p <0.0001, ****; C1+C2+C3+C4 vs Rest, p <0.0001, ****.

Despite its ventralizing capability, knockdown of *bmp16* did not produce an overt dorsalization phenotype. However, given that *bmp16* is co-expressed with additional Bmp ligands in the ventral YSL, we reasoned that the redundant activity of additional Bmp ligands might compensate for its loss. We therefore asked whether knockdown of *bmp16* would sensitize embryos to additional perturbations to overall BMP signaling. Embryos were microinjected control MO, *bmp16* MO, *bmp2b* MO, or both *bmp16* MO and *bmp2b* MO at the one-cell stage and scored for dorsoventral phenotypes at 24 hpf. We identified a dose of *bmp2b* MO that produced a range of dorsalizations such that knockdown of *bmp2b* produced dorsalizations from mild (C1) to severe (C5). *bmp16* knockdown resulted in yolk-specific morphological defects similar to those described above (**Fig. 5A**) (Mullins et al. 1996; Kishimoto et al. 1997). By contrast, combined knockdown of *bmp16* and *bmp2b* resulted in more severe dorsalizations than observed in partial *bmp2b* knockdown alone (**Figs. 5A-B**), demonstrating that these ligands function partially redundantly to promote dorsoventral patterning.

To further characterize *bmp16* and *bmp2b* morphant yolk extension abnormalities, we developed morphological criteria to grade yolk extension phenotypes based on width and length (see Methods). Embryos were classified as mildly narrowed and shortened (YE1; 5-25% decrease in both dimensions), as severely narrowed and mildly shortened (YE2; 25-95% width decrease, 5-25% length decrease), as severely narrowed and shortened (YE3; 25-95% decrease in both dimensions), or as no yolk extension (YE4; 95-100% decrease). *bmp16* knockdown embryos exclusively exhibited YE2 phenotypes, while *bmp2b* knockdown embryos displayed a range spanning YE1 to YE4 (**Figs. S5I-J**). Combined *bmp2b* and *bmp16* MO knockdown produced a shift toward more severe yolk extension defects compared to *bmp2b* MO alone (**Figs. 5SI-J**). Together, these findings indicate that *bmp16* acts in a redundant manner with *bmp2b* to promote dorsoventral patterning and yolk extension morphogenesis.

### YSL *bmp2b* and *bmp16* are maintained independently of BMP autoregulation

Having established that *bmp16* contributes to dorsoventral patterning, we next sought to determine how BMP ligand expression is regulated in the YSL. In the embryo proper during gastrulation, Bmp ligand expression is initially established by early inductive signals and subsequently maintained through BMP-dependent positive feedback (Ramel and Hill 2013). Ventral enrichment of phosphorylated Smad1/5/9 (pSmad) is a well-established readout of BMP pathway activity that has been extensively characterized in embryonic cells throughout gastrulation (J. A. Tucker, Mintzer, and Mullins 2008; Zinski et al. 2017). Pharmacological inhibition of BMP signaling with the type I receptor inhibitor dorsomorphin homolog 1 (DMH1) disrupts this autoregulatory loop, resulting in a loss of Bmp ligand expression (Hao et al. 2010).

However, whether pSmad ventral enrichment extends to extra-embryonic YSL has not been systematically examined. We treated embryos with DMH1 (50 μM) or vehicle control (1.0% DMSO) from 3-8 hpf and immunostained for pSmad. To distinguish between embryonic and YSL nuclei, we developed a segmentation workflow that quantified immunofluorescence (IF) signals from each population independently (see Methods). We confirmed ventral enrichment of pSmad in control treated embryonic nuclei (**Fig. 6A**), and we also detected pSmad-positive ventral YSN (**Figs. 6B, 6SA**). Upon DMH1 treatment, pSmad was abolished in both embryonic cells (as expected) but also in YSN **(Figs. 6A-D, 6SA)**, indicating the YSL requires BMP type I receptor activation to transduce BMP signaling.

**Fig. 6.**
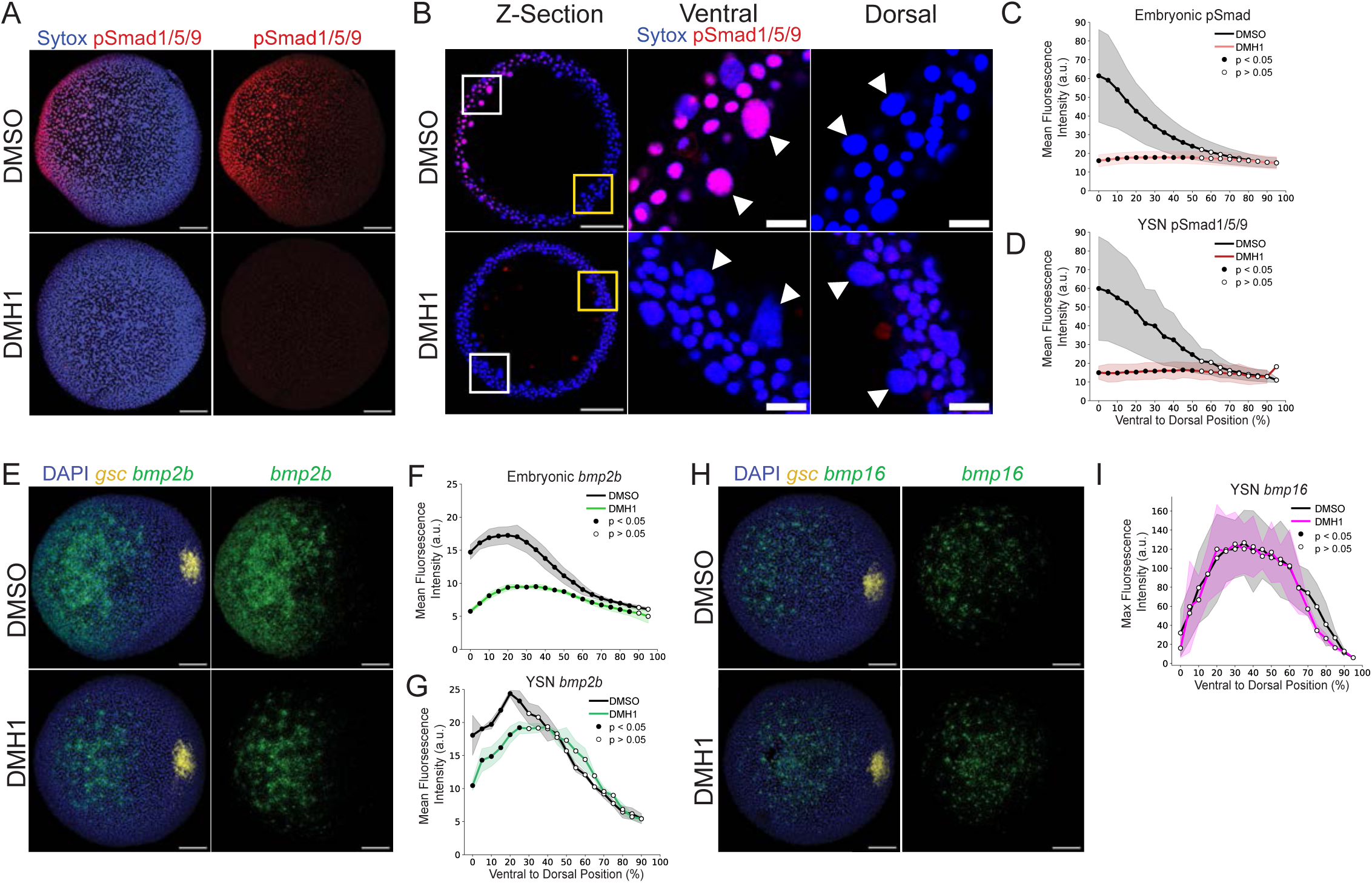
Yolk syncytial *bmp2b* and *bmp16* are maintained independently of BMP autoregulation. **(A)** Representative images of pSmad1/5/9 immunostaining in embryos treated with 1.0% DMSO (top panel) or 50 µM DMH1 (bottom panels) from 3-8 hpf. **(B)** 6 µm Z-section of representative images of pSmad1/5/9 immunostaining treated with 1.0% DMSO (top panel) or 50 µM DMH1 (bottom panels) from 3-8 hpf. Insets from the dorsal (right panels) and ventral (middle panels) axes of embryo with white arrows indicating individual YSN. Scale bar = 20 µm. **(C-D)** Quantification of mean fluorescence intensity of pSmad1/5/9 in embryonic cells (C) and YSN (**D**) across the DV axis in embryos treated with 50 µM DMH1 or 1.0% DMSO from 3-8 hpf. Shaded regions indicate standard deviation. Black dots indicate p > 0.05 and white dots indicate p < 0.05 from Welch’s t-test with Benjamini-Hochberg FDR correction (n = 20 DMSO, 19 DMH1). (**E**) Representative images of *bmp2b* RNA FISH in embryos treated with 1.0% DMSO (top panel) or 50 µM DMH1 (bottom panels) from 3-8 hpf. (**F**-**G**) Quantification of mean fluorescence intensity of *bmp2b* RNA FISH in embryonic cells (**F**) and YSN (**G**) across the DV axis in embryos treated with 50 µM DMH1 or 1.0% DMSO from 3-8 hpf. Shaded regions indicate standard deviation. Black dots indicate p > 0.05 and white dots indicate p < 0.05 from Welch’s t-test with Benjamini-Hochberg FDR correction (n = 19 DMSO, 19 DMH1). (**H**) Representative images of *bmp2b* RNA FISH in embryos treated with 1.0% DMSO (top panels) or 50 µM DMH1 (bottom panels) from 3-8 hpf. **(I)** Quantification of max fluorescence intensity of *bmp16* RNA FISH in YSN across the DV axis in embryos treated with 50 µM DMH1 or 1.0% DMSO from 3-8 hpf. Shaded regions indicate standard deviation. Black dots indicate p > 0.05 and white dots indicate p < 0.05 from Welch’s t-test with Benjamini-Hochberg FDR correction. (n = 24 DMSO, 20 DMH1) *Max projection of the animal view, and dorsal is to the right. Scale bar, 100 µm. Unless otherwise noted. DV, dorsoventral. FDR, false discovery rate*.

We next asked whether expression of Bmp ligands from the YSL is regulated through the canonical BMP-dependent positive feedback loop. Using HCR-FISH, we assessed Bmp ligand expression in both YSN and embryonic nuclei following DMH1 treatment. Upon DMH1 treatment, embryonic *bmp2b* expression was significantly reduced (**Figs. 6E-F**). However, within the YSL, *bmp2b* and *bmp16* expression were each maintained at high levels in both control and DMH1 treated samples (**Figs. 6E, G-I**). Together, these findings indicate that although YSN can transduce BMP signals, the maintenance of YSL *bmp16* and *bmp2b* expression is independent of canonical BMP autoregulation. This indicates that embryonic cells and the YSL maintain expression of *bmp2b* and *bmp16* ligands through distinct regulatory mechanisms.

### Distinct transcriptional networks regulate YSL BMP ligand expression

To explore the basis of this divergent embryonic and extraembryonic BMP regulatory mechanism, we sought to probe the transcriptional regulators governing YSL bmp expression. *mxtx2* is a master regulator of YSL specification, where loss of *mxtx2* results in severe YSL gene misexpression and epiboly failures (Hong et al. 2011; Liu et al. 2024; C. Xu et al. 2012; Bruce et al. 2005; Wilkins et al. 2008). We therefore asked whether Bmp ligand expression within the YSL depends on *mxtx2*. We injected embryos with an *mxtx2* MO at the one-cell stage and performed RNA FISH for Bmp ligand expression. Knockdown of *mxtx2* led to loss of expression of the pan-YSL marker, *apoa1b*, confirming disruption of YSL identity (**Fig. 7SA**). We also found that *bmp2b* expression was decreased in both the YSN and, to a lesser extent, in embryonic cells **(Figs. 7A-C, S7B**). However, *bmp16* expression, which remains YSL-specific at this stage of development, was completely abolished in *mxtx2* morphants (**Fig. 7D-E, 7SC**). These data suggest that YSN expression of *bmp16* and *bmp2b*, are dependent on the YSL transcriptional program established by *mxtx2*.

**Fig. 7.**
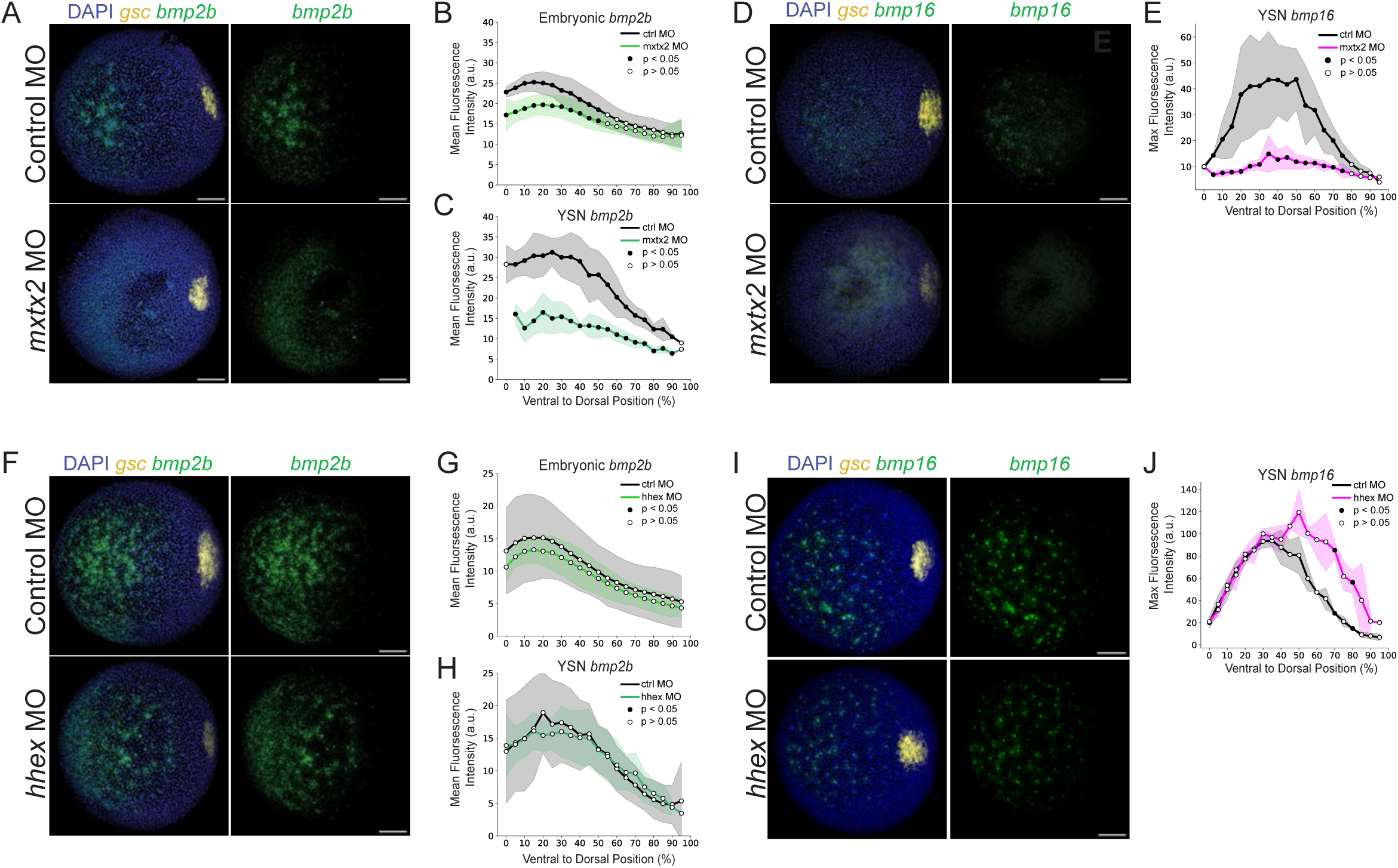
Distinct transcriptional networks regulate YSL BMP ligand expression. (**A**) Representative images of *bmp2b* RNA FISH at 8 hpf of embryos injected at the one- cell stage with Control MO (top panel) or *mxtx2* MO (bottom panels). (**B-C**) Quantification of mean fluorescence intensity of *bmp2b* RNA FISH in embryonic cells (**B**) and YSN (**C**) across the DV axis n embryos treated with 50 µM DMH1 or 1.0% DMSO from 3-8 hpf. Shaded regions indicate standard deviation. Black dots indicate p > 0.05 and white dots indicate p < 0.05 from Welch’s t-test with Benjamini-Hochberg FDR correction (n = 19 control MO, 18 *mxtx2* MO) **(D)** Representative images of *bmp16* RNA FISH at 8 hpf in embryos injected with Control MO (top panel) or *mxtx2* MO (bottom panels at the one-cell stage. **(E)** Quantification of max fluorescence intensity of *bmp16* RNA FISH in YSN across the DV axis at 8 hpf in embryos injected at the one-cell stage with Control MO or *mxtx2* MO. Shaded regions indicate standard deviation. Black dots indicate p > 0.05 and white dots indicate p < 0.05 from Welch’s t-test with Benjamini-Hochberg FDR correction. (n = 15 Control MO, 17 *mxtx2* MO) **(F)** Representative images of *bmp2b* RNA FISH at 8 hpf of embryos injected at the one- cell stage with control MO (top panel) or *hhex* MO (bottom panels). (**G-H**) Quantification of mean fluorescence intensity of *bmp2b* RNA FISH in embryonic cells (**G**) and YSN (**H**) across the DV axis n embryos treated with 50 µM DMH1 or 1.0% DMSO from 3-8 hpf. Shaded regions indicate standard deviation. Black dots indicate p > 0.05 and white dots indicate p < 0.05 from Welch’s t-test with Benjamini-Hochberg FDR correction (n = 18 control MO, 21 *hhex* MO) **(I)** Representative images of *bmp16* RNA FISH at 8 hpf in embryos injected with Control MO (top panel) or *hhex* MO (bottom panels at the one-cell stage. **(J)** Quantification of max fluorescence intensity of *bmp16* RNA FISH in YSN across the DV axis at 8 hpf in embryos injected at the one-cell stage with control MO or *hhex* MO. Shaded regions indicate standard deviation. Black dots indicate p > 0.05 and white dots indicate p < 0.05 from Welch’s t-test with Benjamini-Hochberg FDR correction (n = 20 control MO, 20 *hhex* MO). *Max projection of the animal view, and dorsal is to the right. Scale bar, 100 µm. Unless otherwise noted. DV, dorsoventral. FDR, false discovery rate*.

Having established that *bmp16* expression is downstream of the YSL transcriptional program, we next asked how its expression is restricted to the ventral YSL. Hhex is a YSL-specific transcriptional repressor during dorsal specification of the zebrafish organizer (Bischof and Driever 2004). We asked whether YSL *bmp2b* and *bmp16* ventral expression were maintained by *hhex*. Following *hhex* MO knockdown, we found that both embryonic and YSL *bmp2b* expression patterns remained largely unchanged (**Figs. 7F-H, S7D**). By contrast, *bmp16* expression expanded dorsally in *hhex* morphants (**Figs. 7I-J, S7E**). These findings indicate that *hhex* restricts the *bmp16* expression within the dorsal YSN. Together, these data confirm that YSL *bmp2b* ligand expression is governed by a YSL-specific network rather than canonical BMP autoregulation. While *mxtx2* establishes YSL *bmp16* expression competency, *hhex* spatially refines YSL-specific *bmp16* expression along the dorsoventral axis.

## Discussion

While zebrafish dorsoventral patterning has been well characterized through the lens of a BMP morphogen gradient maintained by embryonic autoregulation, we show that extra-embryonic BMP signaling contributes to dorsoventral axis patterning through a distinct and spatially-regulated extra-embryonic regulatory network. We first established that the YSL is transcriptionally heterogeneous through snRNA-sequencing during gastrulation. Leveraging spatial transcriptomics data, we revealed independent axes of transcriptional covariation that projected onto dorsoventral and animal-vegetal domains of the YSL, ultimately establishing that the YSL is patterned along the same axes as gastrulation stage embryos (Wan et al. 2026). We identified a YSL-specific ventrally expressed gene, *bmp16*, which belongs to the BMP2/4 subfamily and acts redundantly with *bmp2b* to reinforce the BMP morphogen gradient. Notably, we find that the regulatory logic driving YSL BMP expression is distinct from the embryonic BMP autoregulatory pathway. Within the YSL, expression of both *bmp2b* and *bmp16* are independent of BMP pathway autoregulation, demonstrating that the YSL maintains BMP expression through a distinct regulatory program. We establish that *bmp16* and in part, *bmp2b*, expression is dependent on a YSL specific transcription factor, *mxtx2*. We also find that a dorsal YSL determinant, *hhex*, establishes a spatial domain that restricts *bmp16* ventrally. Taken together, we propose a revised framework in which *bmp2b* and *bmp16* ligands are regulated by two parallel pathways including an embryonic regulatory network maintained through BMP autoregulation and an extra-embryonic network maintained through *mxtx2* and *hhex*.

While *bmp16* ventralizes embryos when ectopically expressed, we find that *bmp16* also contributes to endogenous dorsoventral patterning in loss of function contexts. Loss of *bmp16* alone produced a previously undescribed yolk extension phenotype rather than dorsalizing the embryo, while loss of *bmp16* results in dorsalization only when embryos were sensitized by partial loss of *bmp2b*. This pattern is consistent with a model in which *bmp16* and *bmp2b* function in parallel to modulate overall levels of BMP signaling activity. Notably, our spatial projections indicate that while *bmp2b* and *bmp16* YSL expression partially overlap, *bmp2b* spans a larger ventral-animal domain than does *bmp16*. This could suggest that *bmp2b* and *bmp16* are regulated by molecularly distinct, mechanisms and warrants further investigation. Furthermore, the yolk extension phenotype we observed in *bmp16* morphants suggests that *bmp16* fulfills an independent, YSL-specific role. Using a publicly available scRNA-seq atlas data of zebrafish development, we confirm that *bmp16* is expressed in the ventral mesoderm-derived heart and vasculature (**Fig. S3M-N)** (Aponte-Santiago, Su, and Wagner 2026). Interestingly, *bmp16* has been recently demonstrated to regulate cardiovascular morphogenesis (M. Xu et al. 2026). Further dissection of the independent roles played by *bmp16* in extra- embryonic signaling and mesodermal regulation remains of interest.

Our work extends upon previous literature that has identified roles of extra-embryonic tissues in patterning different embryonic axes, with Nodal patterning mesendodermal tissues and BMP patterning ventral tissues (Sun et al. 2014; Chen and Kimelman 2000; Fan et al. 2007). Previous work showed that MGA, Max, and Smad4 act in the YSL to regulate *bmp2b* ligand and are similarly required to specify the ventral tailfin. However, our snRNA-seq data show that Max, Smad4, and MGA are expressed ubiquitously in the YSL, indicating that this regulatory pathway cannot solely account for the ventral spatial restriction of *bmp2b* and *bmp16* (Sun et al. 2014). While we identified *hhex* as an upstream regulator of the spatial patterning of *bmp16* expression, it remains unclear how YSL *bmp2b* expression is spatially restricted. It is possible that *bmp2b* and *bmp16* are restricted to the ventral YSL domain by independent, parallel mechanisms.

While the YSL performs lipid transport and metabolic functions in a role analogous to that of the liver, we find that its transcriptome remains largely distinct, suggesting that these two tissues retain only partially overlapping transcriptional programs. YSL-derived signals also influence liver formation at later stages of development. *rbp4* is a posteriorly expressed YSN- specific gene, and its knockdown causes duplicated liver buds likely due to a liver progenitor migration failure (Li, Korzh, and Gong 2007). Interestingly, *rbp4* is similarly expressed in the visceral extraembryonic endoderm of the rodent yolk sac where it mediates retinol transfer from maternal blood to the fetus, and similar expression has been reported in chick (Soprano, Soprano, and Goodman 1986; Johansson et al. 1997; Yadgary, Wong, and Uni 2014). Extra- embryonic tissues serve critical multifunctional roles to reinforce and actively instruct development of embryonic cells. Whether extra-embryonic spatial transcriptional heterogeneity with functional consequences is conserved across species and inquiry into how syncytial tissues establish and maintain transcriptional heterogeneity across shared cytoplasm remains of interest.

Beyond *bmp16,* the snRNA-seq resource presented here identified many additional genes expressed in a coordinated, spatially organized fashion within the YSL during gastrulation. Additionally, several genes spatially localized within the YSL at later stages of development that have been demonstrated to affect embryonic patterning (Li, Korzh, and Gong 2007; Sakaguchi et al. 2006). While systematic profiling of the YSL throughout development might reveal key signaling genes, it will remain crucial to distinguish YSL-specific roles of genes that are expressed in both embryonic and extra-embryonic tissues. Together, our findings reveal that the YSL is a source of BMP ligands that functions independently from embryonic BMP autoregulation during dorsoventral patterning. This dual BMP regulation could suggest that extraembryonic tissues maintain robustness in signaling pathways during axis specification.

## Supplemental Figures

**Fig. S1.**
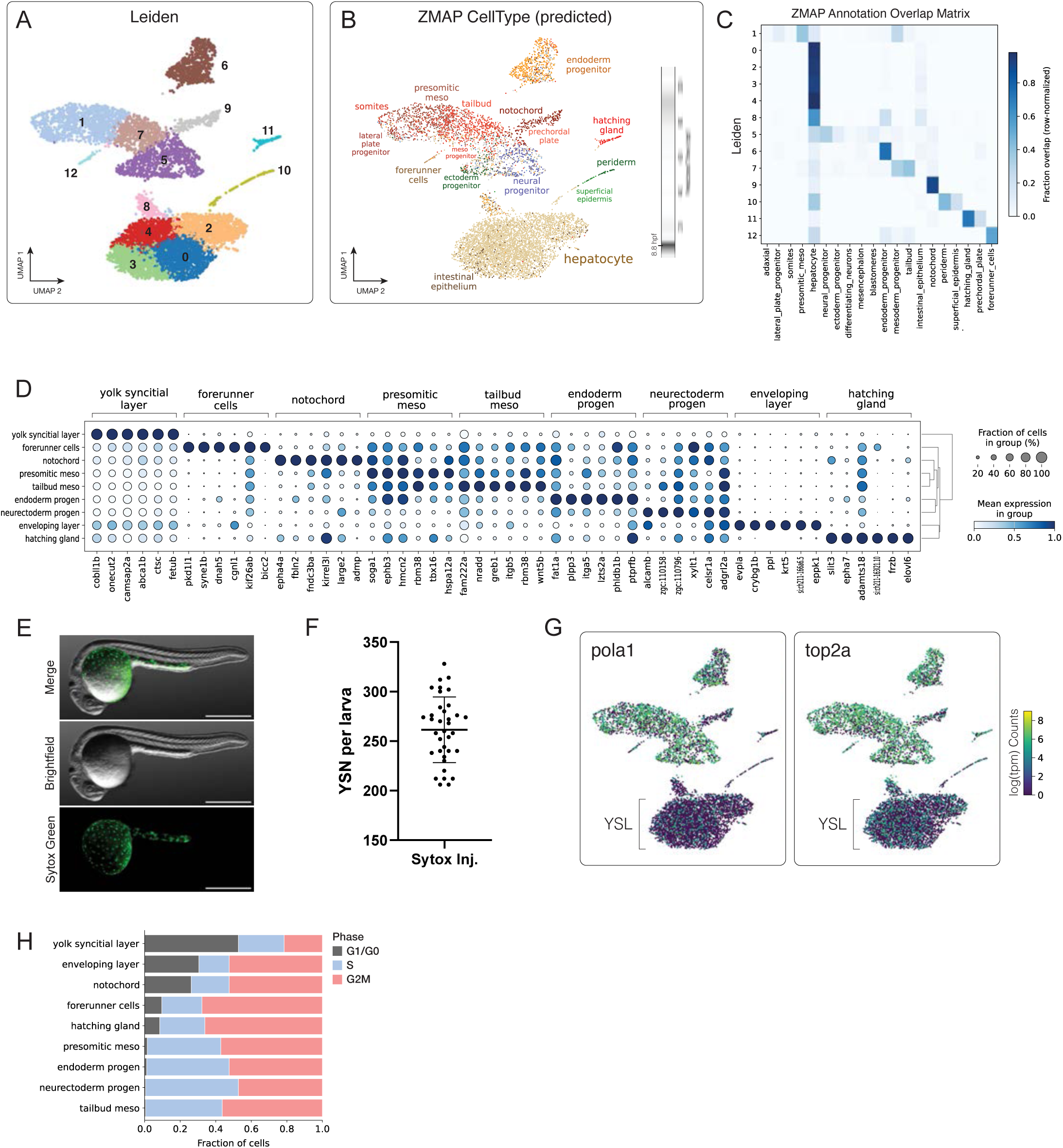
snRNA-seq yolk syncytial nuclei annotation. **(A)** UMAP embedding of all nuclei colored by Leiden cluster assignment. **(B)** UMAP embedding colored by Zebrafish Multi-Atlas Project (ZMAP)-assigned annotations. Histogram of predicted timepoints is depicted as a vertical heat strip. **(C)** Overlap matrix mapping of Leiden clusters to ZMAP annotations. Fractions of cells showing each label pair (row normalized) are indicated via colormap. **(D)** Dotplot analysis of the top 6 enriched transcripts for each annotated cell group. Dot color and size indicate expression level and fraction of expressing nuclei, respectively. **(E)** Representative image of 24 hpf larva injected with SYTOX Green at 3 hpf into the yolk. Scale bar, 500 µm. **(F)** Quantification of number of SYTOX green YSN per larva at 24 hpf. **(G)** UMAP embedding of all nuclei colored by log(tpm) transcript counts for *pola1* (S-phase-associated) and *top2a* (G2M-phase-associated). tpm, transcripts-per-million **(H)** Stacked barplot indicating cell cycle score-derived assignments for G2M-phase, S-phase, and G1/G0 phase for each annotated group.

**Fig. S2.**
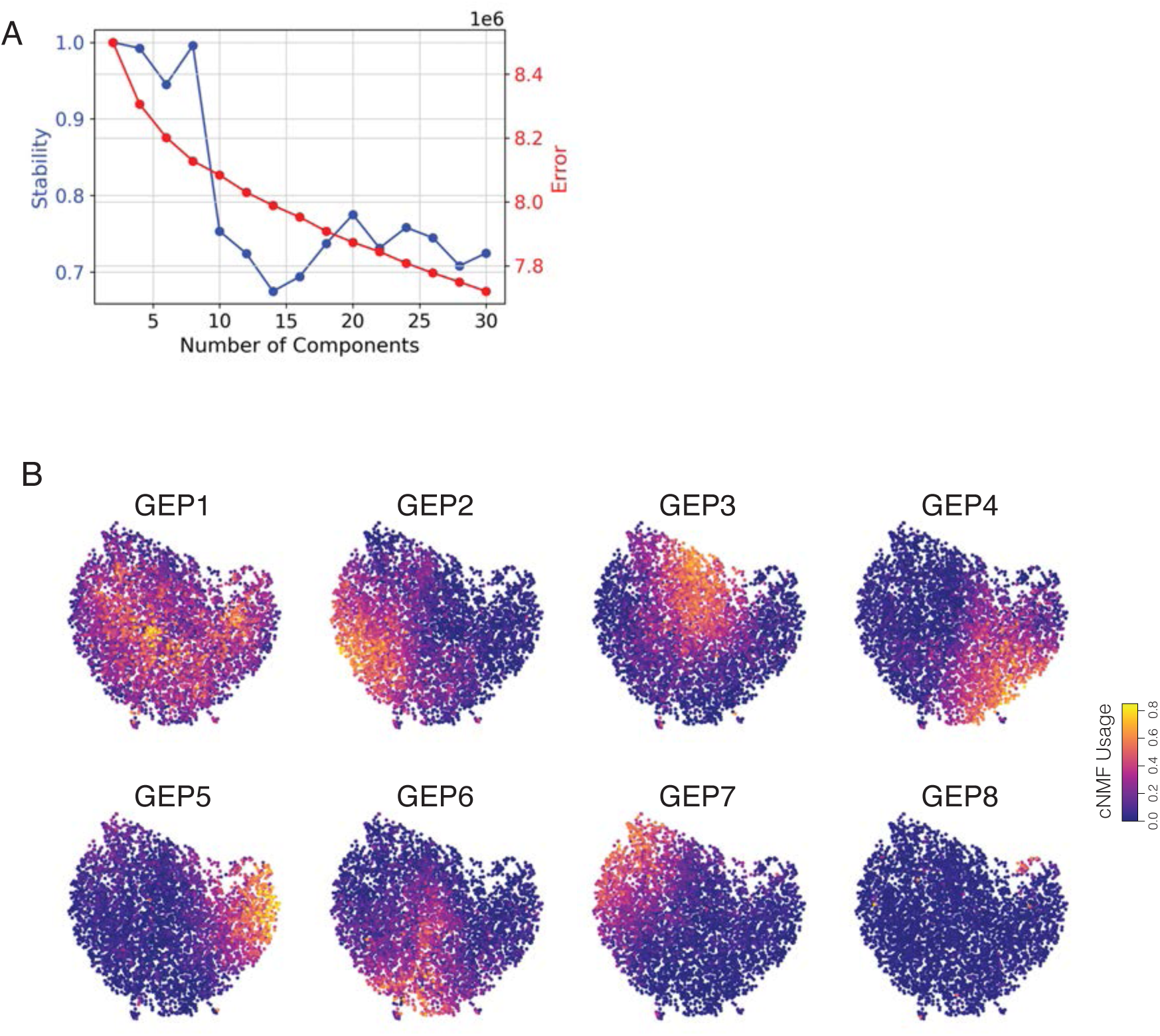
Yolk syncytial nuclei heterogeneity along dorsoventral and animal-vegetal axes. **(A)** Frobenius norm vs stability k-selection plot for YSL subset cNMF. **(B)** UMAP overlays of YSL subset, colored by cNMF GEP scores (8-component factorization).

**Fig. S3.**
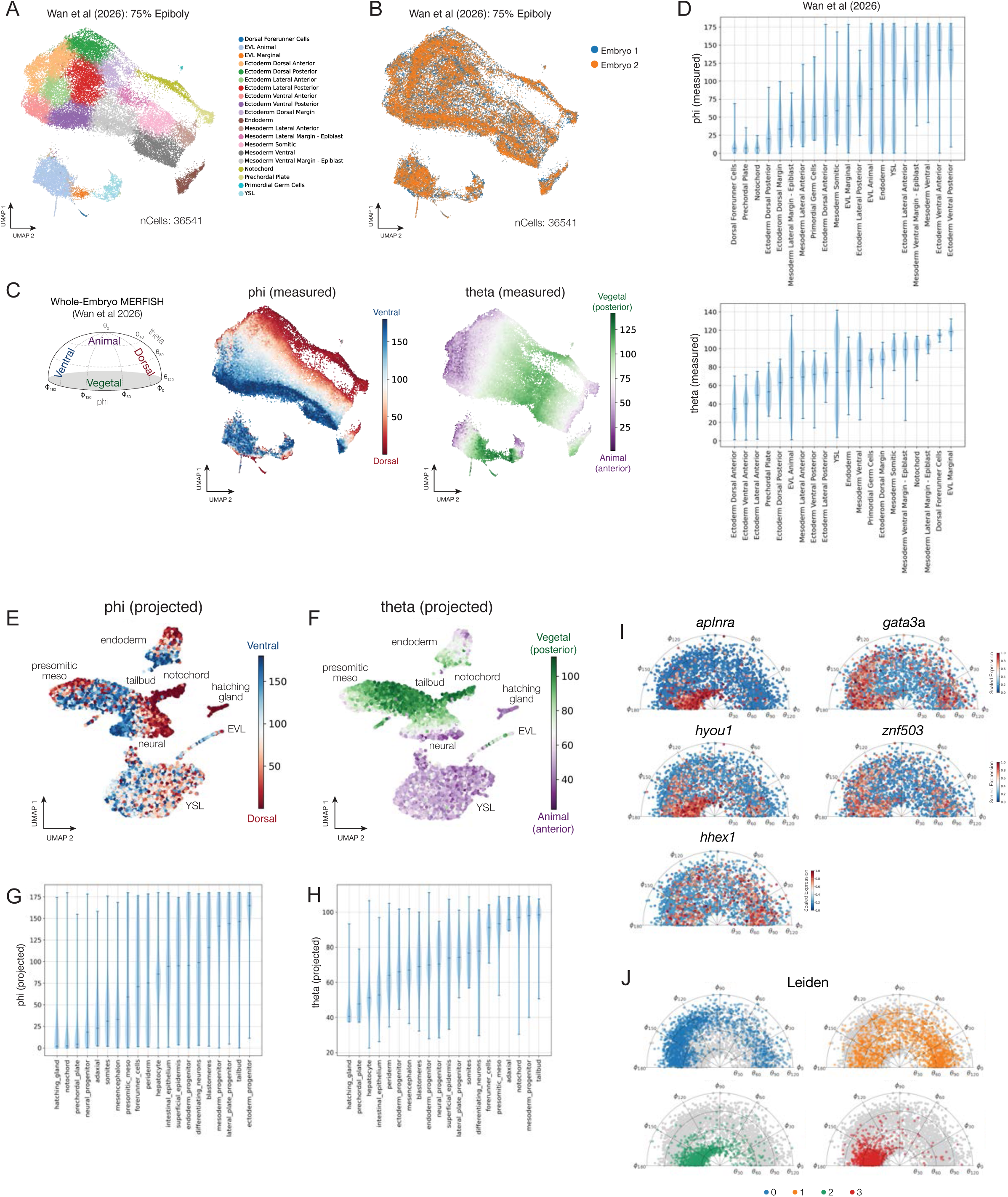
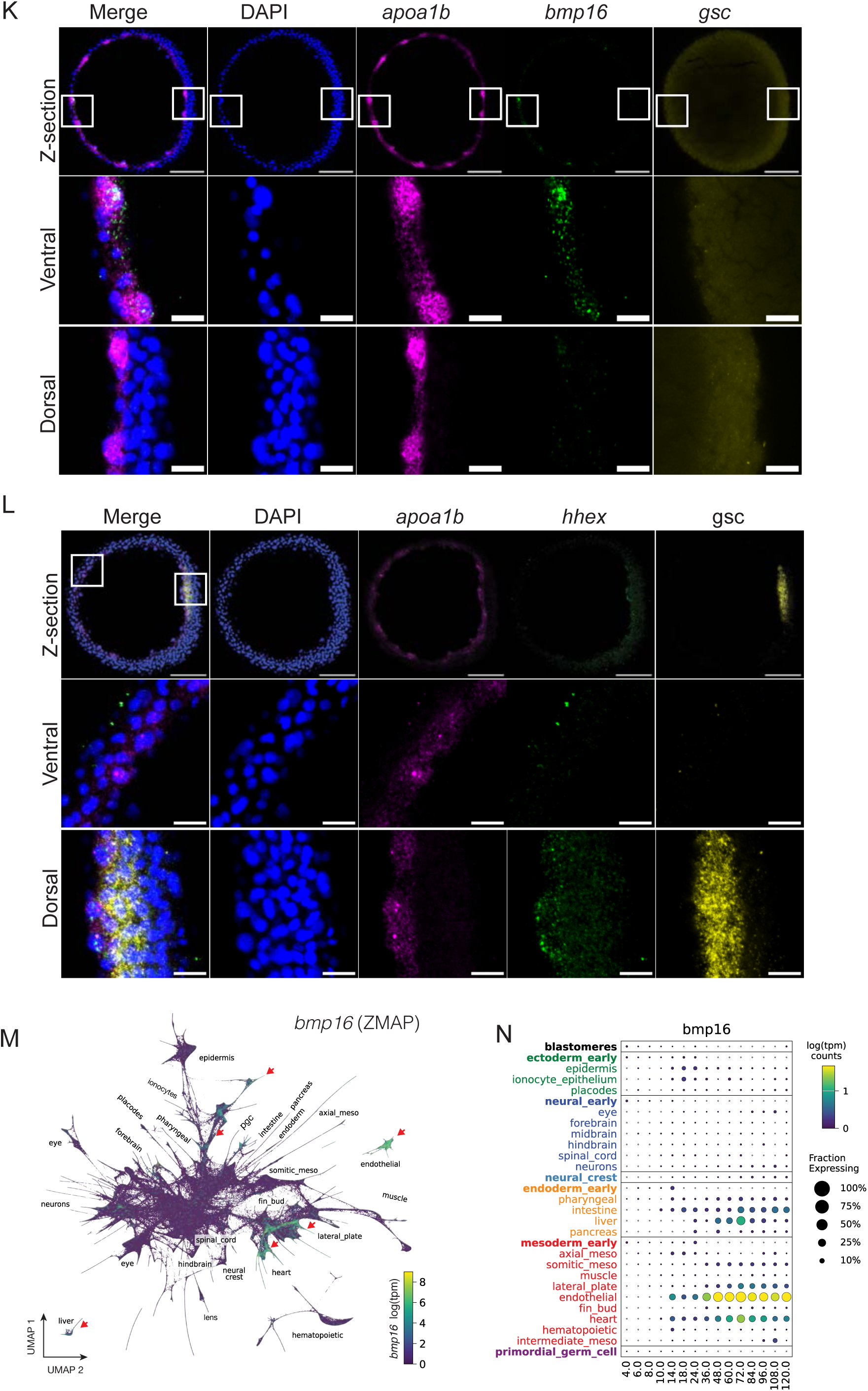
Spatial projections of snRNA-seq data (A-C) Symphony embedding UMAP for (Wan et al. 2026): 75% epiboly dataset. Published cell type annotations (A), embryo of origin (B), and polar coordinates phi and theta (C), are indicated. **(D)** Violin plots for phi (top) and theta (bottom) values for each cell (A-C) grouped by cell type annotation. Polar coordinates stratify cell types by known relative positions. **(E)** phi projections over entire snRNA-seq dataset, select cell type annotations shown. **(F)** theta projections over entire snRNA-seq dataset, select cell type annotations shown. **(G-H)** Violin plots for projected phi (G) and theta (H) values for each cell in (E-F), grouped by annotations. Projected spatial coordinates stratify tissues in accordance with dorsoventral and animal-vegetal localizations. **(I)** YSL polar coordinate projections for select marker genes. Colors indicate scaled relative expression over tpm counts. **(J)** YSL polar coordinate projections for YSL subclusters. **(K)** Representative image of 6 µm Z-stack of *bmp16* RNA FISH. Insets of ventral and dorsal regions (middle and right panels, respectively). Scale bar, 20 µm. **(L)** Representative image of 6 µm Z-stack of *hhex* RNA FISH. Insets of ventral and dorsal regions (middle and right panels, respectively). Scale bar, 20 µm. **(M)** UMAP from ZMAP of *bmp16* expression from 3 to 120 hpf. **(N)** Dotplot from ZMAP of *bmp16* expression from 3 to 120 hpf, rows indicate ZMAP- Tissue level annotation groups.

**Fig. S5.**
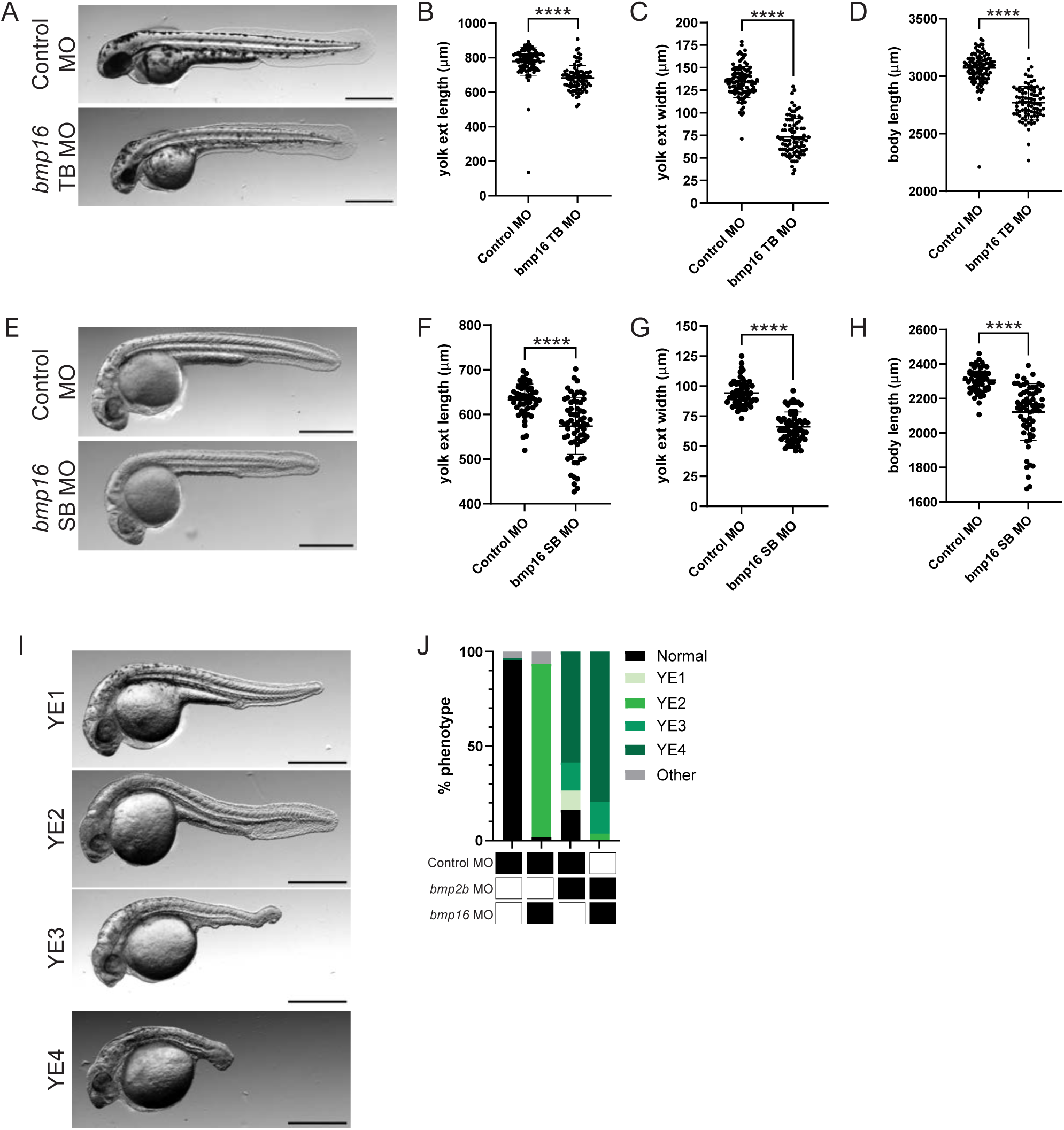
*bmp16* knockdown causes yolk specific defects. **(A)** Representative Images of Control MO (top panel) and *bmp16* translation blocking MO (bottom panel) at 48 hpf. Scale bar, 500 µm. tb, translation blocking **(B-D)** Quantification of *bmp16* translation blocking MO vs Control MO one-cell injection on (B) body axis length, (C) yolk extension width, and (D) yolk extension length at 48 hpf. (n = 104 control MO, 87 *bmp16* MO). Fisher’s Exact test, ****, P < 0.0001. tb, translation blocking **(E)** Representative Images of Control MO (top panel) and *bmp16* splice blocking MO (bottom panel) at 24 hpf. Scale bar, 500 µm. sb, splice blocking **(F-H)** Quantification of *bmp16* TB MO vs Control MO one-cell injection on (B) body axis length, (C) yolk extension width, and (D) yolk extension length at 24 hpf. (n = 52 control MO, 62 *bmp16* SB MO). Fisher’s Exact test, ****, P < 0.0001. sb, splice blocking **(I)** Representative images of yolk extension phenotype abnormalities graded from least severe (YE1) to most severe (YE4) at 24 hpf. **(J)** Stacked bar plot of yolk extension phenotypes in dual *bmp2b* and *bmp16* MO injected embryos. (n = 206 Control MO, 157 *bmp16* MO, 227 *bmp2b* MO, 190 *bmp2b*/*bmp16* dual MO).

**Fig. S6.**
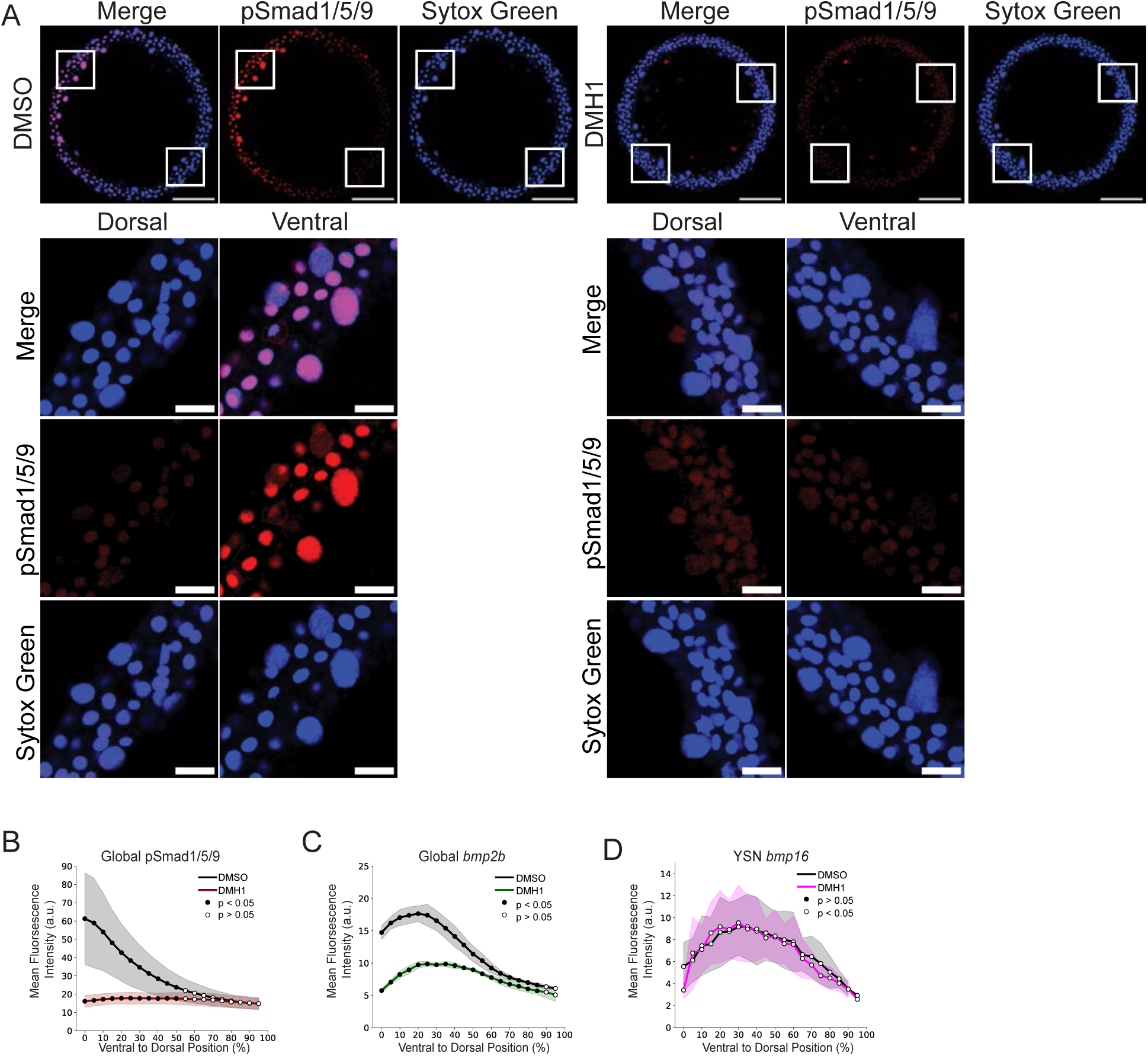
Yolk syncytial nuclei are active in BMP signaling. **(A)** 6 µm Z-section of representative images of pSmad1/5/9 immunostaining treated with 1.0% DMSO (left) or 50 µM DMH1 (right) from 3-8 hpf. Insets from the dorsal (middle panel) and ventral (right panel) axes of embryo show YSN. Scale bar = 20 µm. **(B)** Quantification of mean fluorescence intensity of pSmad1/5/9 in embryonic cells and YSN across the DV axis in embryos treated with 50 µM DMH1 or 1.0% DMSO from 3-8 hpf. Shaded regions indicate standard deviation. Black dots indicate p > 0.05 and white dots indicate p < 0.05 from Welch’s t-test with Benjamini-Hochberg FDR correction. (n = 20 DMSO, 19 DMH1) **(C)** Quantification of mean fluorescence intensity of *bmp2b* RNA FISH in embryonic cells and YSN across the DV axis in embryos treated with 50 µM DMH1 or 1.0% DMSO from 3-8 hpf Shaded regions indicate standard deviation. Black dots indicate p > 0.05 and white dots indicate p < 0.05 from Welch’s t-test with Benjamini-Hochberg FDR correction. (n = 19 DMSO, 19 DMH1). **(D)** Quantification of mean fluorescence intensity of *bmp16* RNA FISH in YSN across the DV axis in embryos treated with 50 µM DMH1 or 1.0% DMSO from 3-8 hpf. Shaded regions indicate standard deviation. Black dots indicate p > 0.05 and white dots indicate p < 0.05 from Welch’s t-test with Benjamini-Hochberg FDR correction. (n = 24 DMSO, 20 DMH1).

**Fig. S7.**
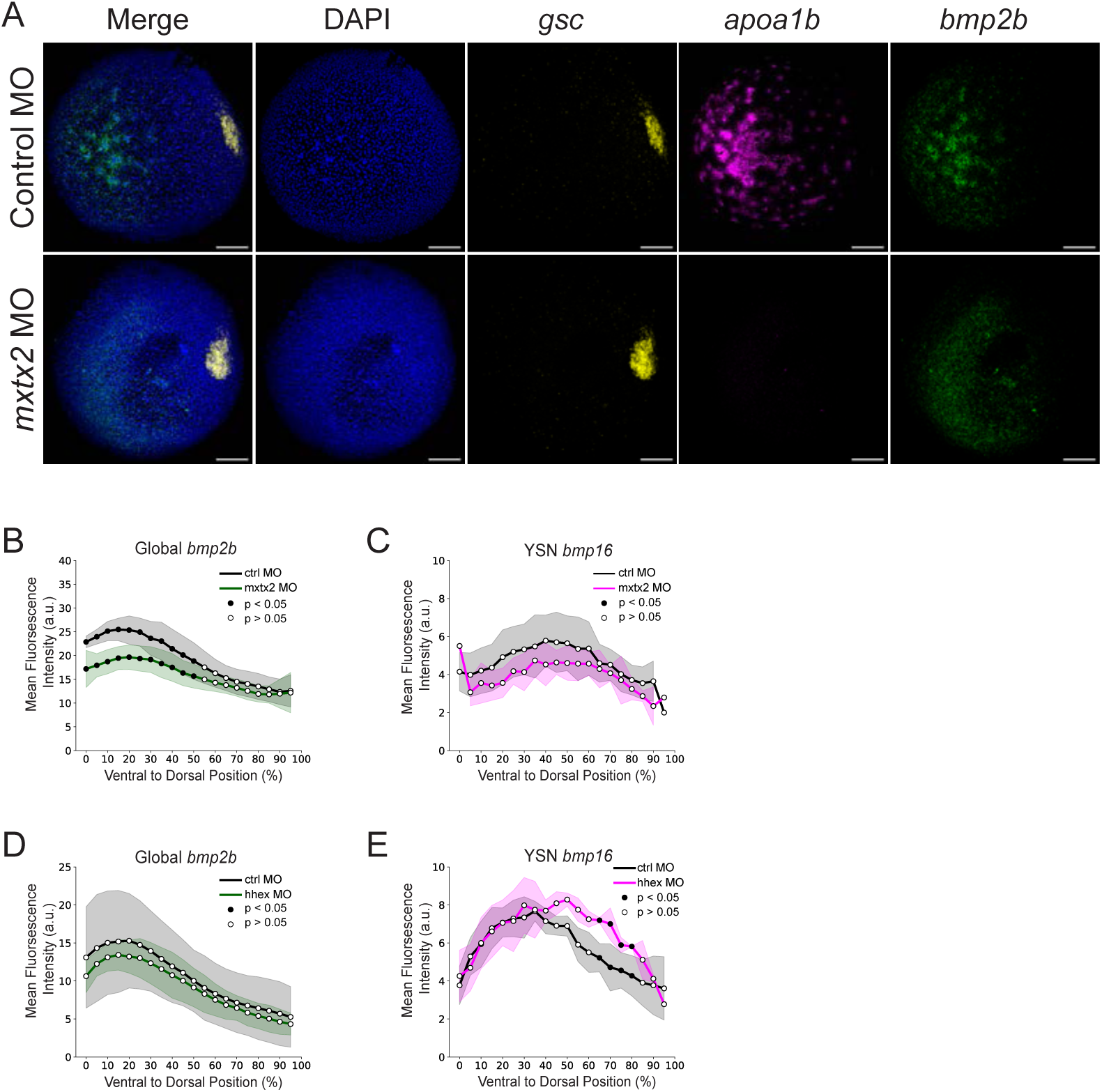
*bmp16* and *bmp2b* regulation through YSL specific regulatory network. **(A)** Representative images of *goosecoid (gsc), apoa1b, and bmp2b* RNA FISH at 8 hpf of embryos injected at the one-cell stage with Control MO (top panel) or *mxtx2* MO (bottom panels). Scale bar, 100 µm. **(B)** Quantification of mean fluorescence intensity of global *bmp2b* RNA FISH in cells and YSN across the DV axis in embryos 8 hpf in embryos injected at the one-cell stage with Control MO or *mxtx2* MO hpf (n = 19 control MO, 18 *mxtx2* MO). **(C)** Quantification of mean fluorescence intensity of *bmp16* RNA FISH in YSN across the DV axis at 8 hpf in embryos injected at the one-cell stage with Control MO or *mxtx2* MO (n = 15 Control MO, 17 *mxtx2* MO). **(D)** Quantification of mean fluorescence intensity of *bmp2b* RNA FISH in embryonic cells and YSN across the DV at 8 hpf in embryos injected at the one-cell stage with control MO or *hhex* MO. Black dots indicate p > 0.05 and white dots indicate p < 0.05 from Welch’s t-test with Benjamini-Hochberg FDR correction. (n = 18 control, 21 *hhex* MO). **(E)** Quantification of mean fluorescence intensity of *bmp16* RNA FISH in YSN across the DV axis at 8 hpf in embryos injected at the one-cell stage with control MO or *hhex* MO. Black dots indicate p > 0.05 and white dots indicate p < 0.05 from Welch’s t-test with Benjamini-Hochberg FDR correction. (n = 20 control MO, 20 *hhex* MO).

## Supplemental Tables

Table S1. Differential expression analysis for YSL-enriched and depleted marker transcripts.

Table S2. GO-TERM enrichment for YSL-associated transcripts

Table S3. YSL subcluster enriched DEGs

Table S4. Gene weights and cell scores for cNMF-defined GEPs

## Materials & Methods

### Zebrafish Husbandry

Adult zebrafish were housed at 28.5°C on a 14-hour light and 10-hour dark cycle. AB wild-type strains were used. Embryos were generated by natural spawning, and the time of fertilization was used to stage each clutch. Later stages were confirmed using morphological criteria.

Embryos were incubated at 28.5°C in 0.3X Danieau medium (10X composition: 174 mM NaCl, 2.1 mM KCl, 1.2 mM MgSO₄, 1.8 mM Ca(NO₃)₂, 15 mM HEPES, pH 7.6) for all wild-type time course experiments and were processed for experiments at the indicated times. All zebrafish experiments were performed in accordance with protocol AN199988 approved by the University of California, San Francisco Institutional Animal Care and Use Committee.

### Microinjections

Morpholinos were by from Gene Tools LLC. Morpholinos were resuspended in water at 100 nmol and stored at room temperature or as aliquots stored at −20°C. Morpholinos were heated to 65°C for 10 minutes before injection and allowed to cool. Prior to injection, all morpholinos were diluted in water and 1X phenol red (Sigma, P0290) to their desired concentration. Embryos were injected at the one-cell stage with 1.0 or 2.0 nL of solution depending on the experimental condition. Morpholinos were injected at the following concentrations mxtx2 (1.0 ng/µL), hhex (1.0 ng/µL), bmp16 SB (0.5 ng/µL), bmp16 TB (2.0 ng/µL), and bmp2b (0.5 ng/µL). To produce a range of milder dorsalizations, 0.25 ng/µL *bmp2b* MO was injected for the *bmp16* mRNA and *bmp2b* MO co-injection experiment. Control MO concentration was adjusted by experimental paradigm to ensure consistent morpholino concentration and volume across experiments.

Sequences for the morpholinos are as follows:

Control MO: CCTCTTACCTCAGTTACAATTTATA

*bmp16* splice blocking MO: TCGGTGAATCTGAATGGAAAAAAGA

*bmp16* translation blocking MO: GAACCAGTAGGCTCGCAGGGAACAT

*bmp2b* MO: CGCGGACCACGGCGACCATGATC

mxtx2 MO: CATTGAGTATTTTGCAGCTCTCTTG

*hhex* MO: GCGCGTGCGGGTGCTGGAATTGCAT

mRNA was resuspended to desired concentration in water and 1X phenol red. mRNA concentration was determined using a dose curve for all mRNAs generated. Stock concentrations were made of mRNAs at 50 ng/µL and then diluted to 5 ng/µL for *bmp2b*, *bmp16*, and mCherry2 mRNA. 1.0 nL of mRNA was injected into the one-cell stage such that each embryo was injected 5 ng of mRNA. For all microinjection experiments with dual injections, separate needles were made for each reagent. Embryos were injected with 1.0 nL solution (2.0 nL total) from 2 different needles.

### mRNA Synthesis

To synthesize *bmp16*, *bmp2b*, and mCherry2 mRNA, we designed and ordered geneblocks from IDT. We included a SP6 and T7 promoter, followed by a Kozak sequence upstream of the coding sequence of each gene based on designs from other expression vectors (Swinburne et al. 2015). An ALFA tag was inserted immediately after the endogenous ATG start site. An Sv40 poly(A)tail and an M13 forward sequence were included on the 3’ end. We PCR amplified the gene block using the primers for Sp6 (ATTTAGGTGACACTATAG) and M13 Forward (GTAAAACGACGGCCAGT) and cleaned up the product with a SPRI bead clean at 0.8X (Beckman Coulter, A63881). In vitro transcription was performed using the T7 promoter and mMessage mRNA transcription kit T7 Transcription Kit according to manufacturer instructions (Thermo Fisher Scientific, AM1344). mRNA concentration was determined using a Nanodrop and correct size confirmed using an Agilent TapeStation 4200 using a High Sensitivity RNA Screen Tape (Agilent, 5067-5579). mRNA aliquots were stored at −80°C. Geneblock sequences are provided in a supplementary table.

### SYTOX Green Injections

A stock concentration of 5mM SYTOX Green (Thermo Fisher Scientific, S7020) was diluted to 50 µM in 0.1 M KCl and injected into the yolk at 3.5 hpf. Injections were deemed successful when YSN only GFP fluorescence was visualized by 4 hpf Leica M205 FCA stereoscope. Because YSN are post-mitotic, larvae were imaged at 24 hpf for YSN counts due to the ease of mounting at this stage relative to 6-8 hpf. In brief, larvae were immobilized with 1X tricaine (Ethyl 3-aminobenzoate methanesulfonate salt; Sigma, A5040) and mounted laterally in an agarose mold. Images were taken on a Leica Thunder. Images were processed on FIJI where each YSN was counted as a ROI. The total number of YSN was multiplied by two as an estimate of total YSN numbers due to the lateral mounting capturing only one side of the larvae.

### Phenotype Characterization

Dorsoventral embryonic phenotypes were characterized at 24 hpf based on well-established criteria. Dorsal phenotypes are characterized as follows: C1 embryos lack part of the ventral tail fin, C2 lack a ventral tail fin or vein, C3 embryos do not have a yolk extension, C4 embryos lack majority of the trunk (fewer than 13 somites), and C5 embryos exhibit radialized dorsal somites notable at 12 hpf and lysis by 24 hpf (Mullins et al. 1996). Ventral phenotypes are characterized as follows: V1 embryos still have a notochord but display anterior neural defects and reduced eye size. V2 embryos have a reduced or lack a notochord, expanded posterior somites with some neural structures. V3 embryos lack have very little or no anterior head structure, no notochord, and enlarged posterior somites. V4 embryos are phenotypically abnormal lacking all anterior structures but still show some posterior somites (Mullins et al. 1996; Kishimoto et al. 1997; Nguyen et al. 1998).

*Bmp16* morpholino phenotypes were characterized based on 3 metrics: body axis length, yolk extension width, and yolk extension length. Yolk extension width was calculated on FIJI (ImageJ) by taking the measurement from a manually drawing a straight line from the top to the bottom of the yolk extension at the center of the yolk extension. Yolk extension length was calculated as the distance from the end of the yolk ball to the urogenital pore opening. Body axis length was calculated from the anterior most region of the head to the posterior most section of the tail. To characterize *bmp16* and *bmp2b* morphant yolk extension abnormalities, we developed criteria to grade yolk extension phenotypes based on its width and length. Embryos were classified as mildly narrowed and shortened (YE1; 5-25% in both dimensions, severely narrowed and mildly shortened (YE2; 25-95% width decrease, 5-25% length decrease, severely narrowed and shortened (YE3; 25-95% decrease in both dimensions), and no yolk extension (YE4; 95-100% decrease).

### Drug Treatments

Dorsomorphin homologue 1 (DMH1; MilliporeSigma, 203646) was resuspended in DMSO (Sigma-Aldrich, 41640) at a stock concentration of 10 mM and stored in aliquots at −80°C C. A working solution of 50 µM was prepared by resuspending the stock solution in the danieau buffer. Embryos were subject to treatment at 28.5°C in a 6 well plate from 3 to 8 hpf.

### Fixation and Permeabilization

Embryos were fixed in 4% paraformaldehyde overnight at 4°C on a rotating carousel. Samples were washed with PBST (0.1% Triton) for 1 hour at room temperature on a rotating carousel. Embryos were manually dechorionated and stored in 0.1% PBST at 4°C for less than one week before downstream processing.

### Immunohistochemistry

Fixed embryos were blocked in FBS-PBSTri (10% fetal bovine serum, 0.1% Triton X-100, 89.9% PBS) for 1 hour at room temperature, followed by incubation in primary antibody (rabbit p-Smad1/5/9 1:100, Cell Signaling, C13820S) diluted in blocking solution overnight at 4°C on a rotating carousel. Each sample was washed 4 x 15 minutes in 0.1% PBSTrit and incubated in fluorescent secondary antibodies overnight at 4°C (goat anti-rabbit IgG Alexa Fluor 647 1:500, Cell Signaling, 4414S). Samples were then washed 4 x 10 minutes in 0.1% PBSTrit. Embryos were then incubated in SYTOX Green (1:2000, Thermo Fisher Scientific, S7020) diluted in PBST for 15 minutes followed by 3 x 10-minute washes in PBST.

### RNA Fluorescence in Situ Hybridization (RNA-FISH)

RNA-FISH was performed using standard reagents and protocols (Molecular Instruments). Embryos were incubated in hybridization buffer for 30 minutes at 37°C and then incubated in probe solution (1-8 pmol of each probe in 250uL probe hybridization buffer) overnight at 37°C. Embryos were washed the following day 4 x 15 minutes in probe wash buffer at 37°C and then washed 3 x 5 minutes in SSCT (3 M NaCl, 0.3 M Sodium Citrate, 0.1% Tween-20, pH 7.0) at room temperature on a rotating carousel. Samples were then briefly washed in 0.1% PBST at room temperature and then incubated in DAPI (1:250) in 0.1% PBST for 1 hour at room temperature on a rotating carousel. Embryos were washed 3 x 10 minutes in 0.1% PBST and then stored overnight at 4°C. Embryos were then mounted in Prolong Gold Antifade Mountant (Thermo Fisher, P10144) for imaging.

### Confocal Imaging and Quantification

To mount samples, fixed samples were deyolked with fine forceps in PBST (0.1% Triton). Samples were mounted on a slide in ProLong Gold Antifade Mountant (Thermo Fisher, P10144) with vegetal pole facing down and animal pole facing up. Vacuum grease was placed around 4 corners of the sample and then a coverslip was placed on top to secure the sample. Samples were immediately imaged after mounting. Images were collected on an upright Leica Stellaris 5 scanning confocal microscope with a 25X Fluotar water immersion lens. Imaris was used to generate 3D renderings of images (Imaris 9.3.1, Oxford Instruments).

Quantification of *bmp2b*/*bmp16* RNA FISH was performed as follows. A reference frame was placed on the widest section of the dorsal side of the embryo based on *gsc* expression, and the diameter from dorsal to ventral was determined by a measurement line manually drawn from gsc across the embryo. All embryonic and yolk syncytial nuclei were segmented on DAPI to determine global signal. This rendering was duplicated and YSN were manually selected based on morphology and *apoa1b* expression to produce 1) embryonic cells only rendering and 2) YSN only rendering. This ultimately produced 3 surface renderings of: (1) global nuclei, (2) embryonic nuclei only, and (3) YSN only. The mean intensity of *bmp2b* or *bmp16* was then calculated for each nucleus across the 3 renderings. The mean intensity was then plotted relative to its position along the X-axis (dorsoventral) relative to *gsc* and binned into 5% intervals along the X-axis (dorsoventral axis).

Quantification of pSmad1/5/9 IHC was performed similarly as above with the following differences. A reference frame was drawn on the dorsal side of the embryo based on pSmad1/5/9 intensity (or morphology of the shield based on thickness of the embryo). A measurement line was manually drawn from dorsal to ventral ends of the embryo to determine the diameter. All embryonic and yolk syncytial nuclei were segmented on SYTOX Green to identify all embryonic and yolk syncytial nuclei. Segmentation was refined as described as above and mean intensity was calculated and plotted the same as above.

Quantification of *bmp16*/*gsc* RNA FISH was performed similarly as described above with the following changes. *Bmp16* mean intensity was calculated based on YSN renderings while *gsc* mean intensity was calculated based on embryonic cell renderings across the DV axis.

*Bmp16* and *gsc* mean intensity were normalized based on the max intensity value in each embryo set to a value of 1. Intensity values were scaled as a proportion of this value and plotted across the dorsoventral axis.

### Single-nuclei dissociations, library preparation, and sequencing

Embryos were manually dissected in the Danieau buffer at 6.5 hpf under a brightfield dissecting scope. Forceps were used to manually remove the blastomere cells from the yolk cell, keeping the yolk cell intact. A 10 µL pipette set to 2 µL was used to pipette the yolk cell into a 1.5 mL Protein Lobind tubes (Eppendorf, 12-698-794) pre-coated in 1.0% Fraction V BSA/DPBS (Gibco, 501215315) kept on ice. 300-600 embryos were dissected over the course of one hour, and then yolks were dissociated by pipetting up and down ten times with a p200 set to 150 uL. Nuclei suspensions were then then fixed using the Parse Biosciences Evercode V2 Nuclei Fixation Kit with minor modifications including: 1) all centrifugation steps done at 10 minutes at 300 g, 2) no filtering was performed, and 3) volume was reduced 4-fold to process samples in 1.5 mL Protein Lobind Tubes instead of 15 mL conical tubes. Fixed sample concentrations were quantified using a hemocytometer and then samples were brought to −80°C in a CoolCell LX cell freezing container (Corning, 432138). Single-nuclei barcoding and library preparation were performed using the Parse Biosciences Evercode WT v2 Barcoding Assay following the manufacturer’s instructions (Parse Biosciences, ECW020301). Final cDNA quality was assessed using an Agilent 4200 Tape Station and Qubit. Samples were submitted to the Chan Zuckerburg Biohub San Francisco for sequencing. 2 final sublibraries were sequenced on an Illumina NextSeq P2 2000 (400 million clusters with the following read configuration: Read 1 (66 bp), Read 2 (86 bp), Index 1 (8 bp), Index 2 (8 bp). Raw FASTQ files were processed using the Parse Biosciences split-pipe pipelines (version 1.0.3p) and reads were aligned to the GRCz11 zebrafish genome assembly. Brief analysis determined insufficient read depth and thus samples were resubmitted for sequencing on an Illumina NovaSeq X 1.5B 2 Lanes (1.5 billion clusters). FASTQ files from both sequencing runs were concatenated and then aligned to the GRCz11 zebrafish genome assembly.

Datasets were analyzed using ScanPy and nuclei were filtered to retain high-quality nuclei based on the following criteria: >700 UMIs, <0.08% mitochondrial gene content, <0.08 doublet score threshold, and <1.0 ribosomal protein mRNA counts. Raw counts for nuclei passing QC filters were normalized by total counts (TPM) and log transformed. The top highly variable genes (n=2000) were identified and scaled for principal component analysis (PCA). PC dimensions (n=22) were estimated based on a permutation based approach comparing PCA eigenvalues to eigenvalues generated from randomized data. Unsupervised leiden clustering was performed with a resolution of 0.6 and the subsequent clustering was annotated using known markers.

### Statistics

Statistical analysis was performed using Graphpad Prism (10.4.2) and Python (3.12.12). All experiments were performed with 3 biological replicates. For image analyses, mean/max fluorescence intensity was calculated for each biological replicate within each distance bin. Differences between control and treated embryos were assessed independently for each distance bin using two-sided Welch’s t-tests. Each embryo was treated as an independent biological replicate. P-values were adjusted for multiple comparisons across distance bins using the Benjamini-Hochberg false discovery rate. Bins with FDR-adjusted p < 0.05 were considered statistically significant. Error bars represent standard standard deviation (SD). We performed a Fisher’s Exact two-sided T-test across dorsalized phenotypes to determine if there was a shift towards more severe dorsalizations. Error bars represent standard error of the mean (SEM). We performed tests across the following bins: C1 vs Rest, C1+C2 vs Rest, C1+C2+C3 vs Rest, and C1+C2+C3+C4 vs Rest. Significance is reported as: P<0.01 *, P<0.001 ** P<0.0001 ***, P<0.00001 ****, and ns = no significance. All other statistical methods are described in the figure legends.

### Spatial Transcriptomics Mapping

To infer embryonic positions of origin for YSL transcriptomes, we transferred spatial coordinates from a whole-embryo MERFISH dataset of two 75% epiboly embryos by k-nearest-neighbor (kNN) projection. Each MERFISH cell is annotated according to spherical coordinates relative to embryonic position: theta (θ), the angular distance from the animal pole, and phi (φ), the angle about the animal-vegetal axis, with phi = 0° at the dorsal midline. To prepare reference data for projections, raw counts were normalized to 10⁴ counts per cell, log-transformed, and z-scored per gene. Principal component analysis (50 components) was performed, and the two embryos were integrated with Harmony (harmony theta = 2). Query scRNA-seq cells went through the same steps: normalization from raw counts, log transformation, restriction to the MERFISH gene panel, and per-gene z-scoring (genes not detected in the query were set to zero). Query cells were then projected into the reference space using the reference PC loadings. For each query cell, the k = 10 nearest reference cells by Euclidean distance were identified. Each neighbor received a Gaussian kernel weight and weights were normalized to sum to 1. theta was assigned as the weighted arithmetic mean of neighbor theta values, using absolute reference values. Because phi represents circular coordinates about the largely symmetrical L/R axis, this coordinate assigned as a weighted circular mean wrapped to (−180°, 180°). This avoids artifact resulting from averaging over the ±180° seam. Left-right asymmetry was therefore collapsed by “folding” phi across the dorsal-ventral axis, giving a dorsoventral coordinate from 0° (dorsal) to 180° (ventral). MERFISH cell-type labels were transferred by the same weighted kNN vote.

Projected cells were plotted in a planar polar layout with the animal pole at the origin. Radial distance equals theta and polar angle equals folded phi. Dorsal (phi = 0°) is to the right, ventral (phi = 180°) to the left, with only the upper half-plane shown (due to phi folding). Cells were colored by Leiden cluster, by projected coordinate, by gene expression, or by cNMF gene expression program (GEP) usage (**Fig. 3B-C, S3I-J**). Each gene’s expression was z-scored across cells and min-max rescaled to [0, 1]. Per-cluster distributions of theta and phi were also shown as violin plots (**Fig. S3D,G,H**).

### ZMAP Annotations

zmap-tools [v0.2.2.2], was used to pre-process, project, and annotate YSL snRNA-seq data using zmap.predict.annotate_with_zmap (**Fig. S1A-C**); annotation was performed at the “CellType” level of the ZMAP ontology and p_thresh was set to 0.5. The ZMAP UMAP embedding and dotplot (**Fig. S3M-N**) were generated using scanpy.pl.embedding and zmap.dotplot.gene_view, using all cells in the ZMAP reference.

### AI Usage

AI-assisted tools were used in the preparation of this manuscript to assist with grammar, clarity, and proofreading of written text. AI-tools were additionally used to support the development of Python-based code. All scientific content, experiment design, data interpretation, and conclusions were generated and verified by the authors.

## Data and Code Availability

Python notebooks used for snRNA-seq data analysis and plotting, as well as ZMAP-based analyses, are available at https://github.com/WagnerLabUCSF/Couturier-2026. Raw and processed data will be made available through the NCBI Gene Expression Omnibus (GEO) and Sequence Read Archive (SRA) repositories upon publication.

## ACKNOWLEDGEMENTS

We thank Y. Su and N. Suren for zebrafish husbandry. We thank C. Chen, C. Chou-Freed, K. Chen, Y. Su, N. Aponte-Santiago, P. Bhat, and I. Roy for discussions and feedback. We thank the Chan Zuckerberg Biohub San Francisco Genomics Platform for genomics support. This work was supported by Chan Zuckerberg Biohub (to D.E.W.), the National Institutes of Health (Grant numbers DP2GM146258 and R00GM121852 to D.E.W.) and the Kinship Foundation Searle Scholar (to D.E.W.).

## Notes

### Competing Interest Statement

The authors have declared no competing interest.

